# A plasma membrane Ca^2+^-dependent protein kinase PtCDPK2 promotes phosphorus starvation resilience in *Phaeodactylum tricornutum*

**DOI:** 10.1101/2025.08.30.671211

**Authors:** Yasmin Meeda, Ellen Harrison, Susan Wharam, Andrea Highfield, Adam Monier, Glen L. Wheeler, Katherine E. Helliwell

## Abstract

Phosphorus (P) is an essential element limiting algal growth and productivity in aquatic ecosystems. Diatoms are important microalgae that thrive in nutrient-variable environments. Determining how diatoms perceive and respond to P availability is therefore crucial for understanding their ecological success. P-limited diatoms use a calcium (Ca^2+^)-dependent signalling pathway to sense and coordinate cellular responses to phosphate resupply. Despite the importance of Ca^2+^ signalling for diatom environmental sensing, apparatus enabling Ca^2+^ signal decoding is poorly understood. Here, we characterise the repertoire of an important group of Ca^2+^ sensor proteins—Ca^2+^ dependent protein kinases (CDPKs), in *Phaeodactylum tricornutum*. Several *PtCDPKs* are transcriptionally upregulated under P starvation. To determine whether PtCDPKs can coordinate P-starvation responses or act to transduce Ca^2+^ signals induced by P resupply, we functionally characterised *PtCDPK2*. PtCDPK2 is highly expressed in P-limited cells and localises to the cell periphery, suggesting a role regulating plasma membrane processes. Further, PtCDPK2 is co-regulated with the transcriptional regulator of P-starvation responses, PtPSR1. PtCDPK2 expression is also coordinated with the induction of P-Ca^2+^ signalling, which is driven by depletion of cellular P rather than external P exhaustion, or growth limitation. *Ptcdpk2* mutants have significantly reduced photosynthetic efficiency and alkaline phosphatase activity under P starvation, but we do not find evidence for a direct role coordinating downstream responses to P resupply. These findings suggest PtCDPK2 is essential for regulating P-starvation physiology and reveals a role for Ca^2+^-signalling apparatus in promoting diatom tolerance in low P environments.

## Introduction

Phosphorus (P) is a crucial macronutrient sustaining life in aquatic environments. P has vital roles as a constituent of key molecules including nucleic acids, phospholipids and ATP, alongside mediating cell signalling (Bünemann and Condron 2007; Paytan and McLaughlin 2007). P limits algal growth in lakes, open ocean and coastal ecosystems (Thingstad et al. 1998; Hudson et al. 2000; Ly et al. 2014). Moreover, changing anthropogenic nutrient inputs are exacerbating P limitation in coastal habitats (Seitzinger et al. 2010; Burson et al. 2016). Yet, P availability can fluctuate considerably due to river run-off, atmospheric deposition and nutrient up-welling (Jordan and Joint 1998; Karl 2014; Duhamel 2024). Mechanisms enabling algae to cope with varying P supply thus govern their competitive success in aquatic environments. We previously discovered that under P-limiting conditions a globally important group of microalgae, the diatoms, detect phosphate resupply via a rapid Ca²⁺ signalling response (Helliwell et al., 2021a). Apparently absent in green algae (*Chlamydomonas reinhardtii*) and plants (*Arabidopsis thaliana*) (Matthus et al. 2019; Pivato et al. 2025), this novel sensory pathway enables diatoms to quickly detect and coordinate recovery from P starvation. However, the molecular machinery underlying P-Ca^2+^ signalling, including how signals are generated and decoded remains unknown.

Diatoms (stramenopiles) contribute approximately 40% of marine primary productivity (Armbrust 2009; Malviya et al. 2016), and are emerging as powerful model systems for algal cell biology (Armbrust et al. 2004; Bowler et al. 2008; Nymark et al. 2016; Mock et al. 2017; Falciatore et al. 2020; Villar et al. 2025). Genomic and genetic resources have yielded important insights of how diatoms cope with P starvation. Diatoms induce phosphate transporters and alkaline phosphatases to maximise P acquisition (Dyhrman et al. 2012; Lin et al. 2013; Cruz de Carvalho et al. 2016; Alipanah et al. 2018; Matsui et al. 2024). Intracellular stores of polyphosphate (PolyP) are also consumed under P deficiency (Lapointe et al. 2024), and cellular usage is economised, including through substitution of phospholipids with alternative forms (e.g. sulfolipids) (Van Mooy et al. 2009; Martin et al. 2011). Additionally, P-starved benthic diatoms glide towards P hotspots (Bondoc et al. 2019). P limitation also causes reductions in nitrogen (N) uptake and photosynthetic efficiency (Cruz de Carvalho et al. 2016; Helliwell et al. 2021a; You et al. 2022).

Despite belonging to a different eukaryotic supergroup compared to green algae and plants (Dorrell and Smith 2011), diatoms share key P-starvation regulatory machinery. The MYB-like transcription factor phosphate starvation regulator 1 (PSR1), first identified in *C. reinhardtii* (Wykoff et al. 1999) and subsequently *A. thaliana* (known as PHR1) (Rubio et al. 2001), is conserved in diverse algae including diatoms (Ngan et al. 2015; Sharma et al. 2020; Fiore et al. 2021). P-starved *C. reinhardtii Crpsr1* mutants have differential gene expression compared to wildtype (Moseley et al. 2006). However, overlap between P-responsive target genes of PSR1/PHR1 is limited between *C. reinhardtii* and *A. thaliana* (Hammond et al. 2003; Wu et al. 2003; Moseley et al. 2006). In the model pennate diatom *Phaeodactylum tricornutum, Ptpsr1* mutants show decreased induction of alkaline phosphatase activity and reduced phospholipid degradation (Sharma et al. 2020). Rapid elevations in cytosolic Ca^2+^ seen in response to phosphate resupply to P starved cells are also impaired (Harrison et al. 2025). This suggests direct or indirect regulation of P-Ca^2+^ signalling machinery by PtPSR1. Pharmacological inhibition of P-Ca^2+^ signals reduces the ability of cells to mediate rapid enhancements in N uptake capacity necessary for recovery from P deficiency (Helliwell et al. 2021a). However, how P-Ca^2+^ signals are transduced to coordinate such downstream responses are completely unknown.

Like other eukaryotes, diatoms use Ca^2+^ signalling to perceive a diverse array of stimuli including mechanical stress, nutrients (phosphate and iron), allelochemicals, temperature, light, membrane depolarisation and osmotic shock (Falciatore et al. 2000; Vardi et al. 2006; Helliwell et al. 2019, 2021b; Kleiner et al. 2022; Flori et al. 2024). Components of the diatom Ca^2+^ toolkit have been deduced, including ion channel repertoire, and functional characterisation of novel single-domain voltage-gated Ca^2+^ channels (Verret et al. 2010; Helliwell et al. 2019; Murphy et al. 2024), as well as Ca^2+^ efflux proteins (Liu et al. 2024). By contrast, diatom Ca^2+^ sensor apparatus has received little attention. Ca^2+^ sensor kinases can detect specific Ca^2+^ signals (or signatures) arising from external stimuli, to induce downstream responses via phosphorylation (Harper et al. 2004; Dodd et al. 2010). Eukaryotic genomes encode distinct families of Ca^2+^ sensors (Bredow and Monaghan 2019). A major family in plants, red and green algae and protists are the Ca^2+^-dependent protein kinases (CDPKs), or CPKs in plants (Valmonte et al.; Billker et al. 2009; Brawley et al. 2017; Bredow and Monaghan 2022). CDPKs contain a calmodulin-like domain typically with four Ca^2+^-binding EF hands, fused to a C-terminal Ser/Thr kinase domain (Harper et al. 1991; Wernimont et al. 2010). They are crucial in transducing nitrate-induced Ca^2+^ signals in *A. thaliana*, orchestrating primary nitrate responses via phosphorylation of NIN-LIKE PROTEIN (NLP) transcription factors (Riveras et al. 2015; Liu et al. 2020). CDPKs also regulate ammonium transporters (Valmonte et al. 2014; Qin et al. 2020), and are upregulated by N and P deprivation in green algae (Motiwalla et al. 2014; Caló et al. 2017).

Whilst CDPKs have received little attention in diatoms, several predicted CDPKs are upregulated under P deprivation in *P. tricornutum* (Cruz de Carvalho et al. 2016). As they contain PtPSR1 motifs in their promoter regions, they are likely controlled by this transcription factor (Sharma et al. 2020). Upregulation of CDPKs during P deficiency suggests they may be required to detect P resupply induced Ca^2+^ elevations. Alternatively, CDPKs could coordinate adaptations to P limitation. To investigate these scenarios, we examined *P. tricornutum* CDPKs, including localisation, expression and functional roles of PtCDPK2 in response to P limitation and resupply, and in relation to P-Ca^2+^ signalling and PtPSR1.

## Methods

### Strains and culturing

The background strain used in this study was *Phaeodactylum tricornutum* CCAP 1055/1. This strain was used to generate the PtCDPK2-mVenus lines, as well as the *Ptcdpk2* mutants and their subsequent derivative lines expressing the genetically encoded Ca^2+^ biosensor R-GECO1-mTurquoise (RGMT) (Harrison et al. 2025), as described below. Additionally, the Ca^2+^-signalling experiments shown in Figure 3, were conducted with a genetically modified strain of *P. tricornutum* CCAP 1055/1 (PtR1), expressing R-GECO1 as previously generated by (Helliwell et al. 2019). The PtPSR1-mVenus line was described in (Harrison et al. 2025). Cultures were grown in filtered seawater (FSW) with f/2 nutrients (phosphate, trace metals and nitrate (Guillard, R.R.L. and Ryther, (1962)) at 18°C under a 16:18 light: dark cycle with a light intensity of 60-80 µE m^-2^ s, unless otherwise stated.

All physiology experiments were inoculated to a starting density of 3 x 10^4^ cells/ml into high (36 µM initial phosphate), low (1.8 µM initial phosphate) or phosphate resupply (inoculated into 1.8 µM initial phosphate and then resupplied with 36 µM phosphate on day 4) treatments. Cell counts of *P. tricornutum* were performed at the same time each day using a Beckman Coulter Counter (Beckman Coulter, UK Ltd, Buckinghamshire, UK) with the aperture set to measure between 3-9 µm. The Coulter Counter was washed through with Isoton (Beckman Coulter, UK) and cell counts commenced once the blank measurement was below 50 cells/ml. Growth rates were calculated using the following equation: μ =ln(N2/N1)/(t2 −t1), where μ is the specific growth rate, and N1 and N2 are the cell counts at time 1 (t1) and time 2 (t2), respectively.

### Photosynthetic efficiency (F_v_/F_m_) measurements

Photosynthetic efficiency Fv/Fm (maximum quantum yield of photosystem II) was measured in triplicate with cells dark-adapted for 20 mins prior to analysis. Measurements were taken using the PAM fluorometer (Water-PAM, Walz, Germany) with 3 ml cuvettes. The gain was set at 5 for *P. tricornutum* reads.

### Identification of *PtCDPK* genes in *P. tricornutum*

To identify CDPKs in the *P. tricornutum* CCAP 1055/1 genome (v3) (Bowler et al. 2008; Villar et al. 2025), an initial sequence similarity search using *A. thaliana* CPK1 (Uniprot Q06850) (Harper et al. 1993, 1994) as a query sequence was performed using BLASTp against the DiatomicBase v3 genome portal using default parameters (Villar et al. 2025). *P. tricornutum* PtCDPK2 (protein id. 21006) was used subsequently to further examine the presence of CDPKs in the *P. tricornutum* genome. Nine CDPK-like hits containing both EF hand (IPR002048) domain/s and a kinase (IPR000719) domain were verified using Interpro analysis (Apweiler et al. 2000), and are presented in **Figure1A**.

**Figure 1.**
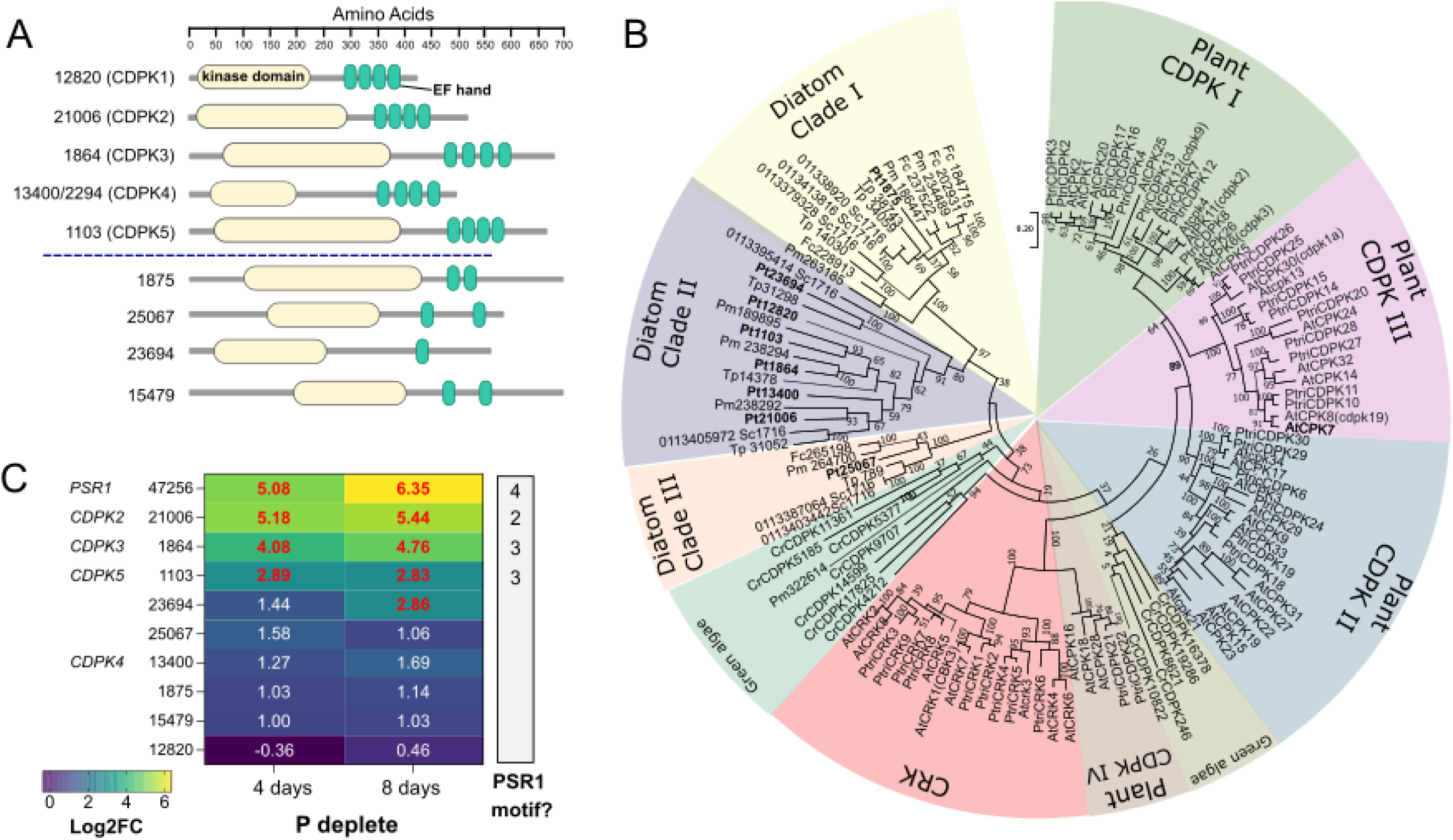
*In silico* analysis of *Phaeodactylum tricornutum* Ca^2+^-dependent protein kinases (CDPKs). **A)** Domain analysis of CDPK-like hits in *P. tricornutum*. Kinase domain in yellow and EF-hands in green. Grey line represents the length of the protein. Protein identifiers for each protein are given. Domain analysis was performed using Interpro (Apweiler et al., 2000). As Pt12820 was previously annotated in the genome as PtCDPK1, Phatr2_21006 is referred to as PtCDPK2 in this study. Above the dotted line shows CDPKs with four EF hands and below the line shows CDPK-like proteins with fewer than four EF hands detected. **B)** A maximum likelihood phylogenetic tree of CDPKs from *P. tricornutum (Pt), A. thaliana (At), Populus trichocarpa (Ptri), Chlamydomonas reinhardtii (Cr), Fragilariopsis cylindrus (Fc), Pseudo-nitzschia multiseries (Pm) Thalassiosira pseudonana (Tp).* The tree was constructed using a WAG+ Freqs (+F) correction model with 100 non-parametric bootstrap replicates, with bootstrap support indicated on the tree. **C)** Heatmap showing log2fold changes in transcriptional expression of *P. tricornutum CDPK* hits during phosphate limitation for 4- and 8-days compared to the 4-day old replete conditions, plotted from a previously published transcriptomic study (Cruz de Carvalho et al., 2016). Genes showing a significant greater than two-fold change (p-value<0.05) in P-deplete conditions are highlighted in red. The number of PtPSR1 recognition motif/s in the promoter region as reported by (Kumar Sharma et al., 2020)is also indicated.

### Phylogenetic analysis

For the reconstruction of the phylogenetic tree, protein sequences from *C. reinhardtii* were obtained from (Li et al. 2019) and plant sequences taken from Xiao et al., (2016); Yip Delormel & Boudsocq, (2019) and Bredow & Monaghan, (2022). In addition to *P. tricornutum* CDPKs, CDPK hits from the diatoms *Pseudo-nitzschia multiseries* CLN-47 (genome v1) and *T. pseudonana* CCMP 1335 (genome v3) were also included, obtained through protein sequence similarity searches of genomes provided via the JGI portal. A maximum likelihood phylogenetic tree was constructed using the protein sequences. Protein amino acid sequences were aligned using MAFFT v7 webserver, using default parameters (Kuraku et al. 2013; Katoh et al. 2019). The final alignment was 464 amino acids in length. Molecular Evolutionary Genetics Analysis (MEGAX) software was used to reconstruct the tree (Kumar et al., 2018) and identify the best-fitting evolutionary model using corrected Akaike Information Criterion. A maximum likelihood tree was generated using the WAG+F model with 100 non-parametric bootstrap replicates.

### PtCDPK2-mVenus construct design and strain generation

To generate the PtCDPK2-mVenus construct a 2604 bp sequence for gene id. 21006, was synthesised and custom cloned by Genscript (GenScript, Piscataway, NJ) using PstI and StuI restrictions sites, into the pPha-T1-mVenus vector (a derivative of the pPHA-T1 (accession AF219942) adapted as described by (Helliwell et al. 2019) containing a codon optimised mVenus gene (AJN91098.1). The synthesised sequences include the start codon (ATG) as well as 700 bp upstream, alongside the gene sequence including exons and introns, but excluding the stop codon (**Supplementary Information 1**). The resulting plasmid was transformed into *P. tricornutum* via biolistic transformation with zeocin selection (75 µg/ml) as outlined previously (Helliwell et al. 2019). Transformants were screened for mVenus fluorescence using the Leica SP8 confocal microscope with a 63x oil objective. Cells were excited with a 488 nm argon/krypton laser and emission was detected between 500-530 for mVenus and 650-710 nm for chlorophyll autofluorescence. For FM4-64 staining, PtCDPK2-mVenus cells were grown in f/2 medium for four days, before addition of 10 μM FM4-64 and incubation for 20 mins at room temperature before imaging. Cells were visualised with a 488-nm laser, and the emission signals were collected by detection windows ranging from 606 to 650 nm on a Leica SP8 confocal microscope.

### Ca^2+^-imaging experiments to define thresholds of P depletion to activate P-Ca^2+^ signalling response

*P. tricornutum* strain PtR1 (Helliwell et al. 2021a) was grown in artificial seawater (ASW) using Tropic Marin Classic (Tropic Marin®) sea salts (30 g/l) with pH adjusted to 8.1 and sterile filtered, with f/2 nutrients added as described before, except for phosphate that was provided at the different concentrations indicated in the results. Glass bottomed dishes 35 mm diameter (In Vitro Scientific, Sunnyvale, CA, USA) were coated with 0.01% poly-L-lysine (Sigma-Aldrich, USA) and subsequently washed with 1 ml ASW without nutrients and PtR1 cells (on day 4 and day 5) were added and left for 10-30 min to settle. Cells were imaged at 20°C using a Leica DMi8 microscope (Leica, Germany) 63 x oil immersion objective. R-GECO1 cells were excited with 530-555nm excitation and emission measured at 575-630 nm, using a SpectraX LED light source (Lumencor). Image capture and analysis were carried out on Leica Application Suite X. Cells were perfused with ASW for 30 s using a gravity-fed perfusion system, then for phosphate resupply treatments, cells were perfused with ASW with 36 μM phosphate (NaH_2_PO4) for 30 secs, unless stated otherwise.

### Generation of *P. tricornutum Ptcdpk2* mutants

The CRISPR-Cas9 vector developed by Nymark et al., (2016) was employed for editing the *PtCDPK2* gene (protein id: 21006) in *P. tricornutum* (Nymark et al. 2016). We designed two sgRNAs targeted to generate an ∼342 bp deletion. This approach has proven efficient previously for high-throughput screening for bi-allelic mutants via PCR in diatoms (Hopes et al. 2016; Helliwell et al. 2019). A library of candidate sgRNAs was generated using the PHYTOCRISPEX (Rastogi et al. 2016) web tool with default parameters (NGG PAM, and CRISPR start from ‘G’). The Broad Institute sgRNA design program (Doench et al. 2016; Sanson et al. 2018) was used subsequently to obtain ‘on-target’ efficiency scores. Two 20 bp guide RNAs (sgRNA1_Pt21006: GAAACAGAACAACAAAAGGA and sgRNA2_Pt21006:

GAAGTATTGGAGTTGTGTGA) that passed the PHYTOCRISPEX OFF-target criteria were chosen based on their ON-target scores (0.57 and 0.66, respectively) and position within the gene. Target sgRNAs were predicted to disrupt the region encoding the kinase domain of the protein. A 343 bp DNA fragment was synthesised (Genscript) containing two tandem U6Promoter:sgRNA:gRNAscaffold:terminator cassettes for sgRNA1_Pt21006 and sgRNA2_Pt21006 respectively (**Supplementary Information 2**) (Hopes et al. 2016). BsaI restriction site overhangs were also included, to clone the DNA fragment into the pKS diaCas9_sgRNA (Addgene: 74923) plasmid. The resulting plasmid was co-transformed along with pPHAT1 (accession no.: AF219942) conferring zeocin resistance using biolistic transformation and 75 µg/ml zeocin for selection, as previously described by (Helliwell et al. 2019). Putative *Ptcdpk2* mutant colonies were screened via PCR using the Phire Plant Direct PCR Kit (ThermoFisher Scientific) with primers targeting the *PtCDPK2* gene: KEH_409F: CGTCCAAGTGCTCGGTAATG and KEH_410R: GTCTCATCTGGCAAGCGTTC. Mutants were further purified by diluting primary colonies suspended in FSW and plating on selection plates (50% FSW f/2 with 75 µg/ml zeocin). Resulting colonies were picked and re-screened via PCR. DNA sequencing produced clean (unmixed) chromatograms were obtained for *Ptcdpk2.1*, *Ptcdpk2.3, Ptcdpk2.4*, and *Ptcdpk2.5,* with sequencing information provided in **Supplementary Information 3-6**. Coding sequences of mutant and WT lines were further analysed via reverse transcriptase (RT)-PCR (**Supplementary Methods**).

### Generation and Ca^2+^-imaging of *Ptcdpk2* mutants expressing R-GECO1-mTurquoise

The WT RGMT strain was as generated and described by (Harrison et al. 2025). *Ptcdpk2.1*, *Ptcdpk2.3*, and *Ptcdpk2.4* mutants were transformed with the RGECO1-mTurquoise_pPHAT1 via biolistic transformation with Blasticidin (8 µg/ml) selection, using the methods outlined previously (Helliwell et al. 2019). Cells were imaged using a Leica DMi8 inverted microscope (Leica Microsystems) with a 63x 1.40 NA oil immersion objective. A SpectraX LED light source (Lumencor) was used with a 550/15 nm excitation filter and 585/40 nm emission for R-GECO1 and a 470/24 nm excitation filter and 525/50 nm emission filter for the mTurquoise, in a sequential manner. As previously detailed, a gravity-fed perfusion system was used, with FSW perfused for 30 secs before switching to FSW + phosphate (36 µM). As the RGMT line can be used as a ratiometric Ca^2+^ indicator, the initial background ROI fluorescence was removed from each channel then the R-GECO1/mTurquoise ratio calculated, this was then normalised by using the ratio of the 6 frames preceding the phosphate resupply stimulus (R/R_0_).

### Quantification of alkaline phosphatase activity

Alkaline phosphatase (AP) activity was quantified using the substrate p-nitrophenyl phosphate (pNPP) (Sigma-Aldrich, N7653) assay, which is based on the cleavage of pNPP by alkaline phosphatase resulting in a yellow substance. The protocol was modified from Shemi et al., (2016). Briefly, 1.5 ml of cells were centrifuged 17,000 g for 5 min and resuspended in a total of 176.9 μl of alkaline phosphatase buffer (0.01 M Tris pH 8, 0.05 M MgCl_2_ and 0.01 M CaCl_2_) with 23.1 µl of pNPP liquid substrate. Absorbance at 405 nm was measured at 2 min intervals during 30 min using a CLARIOstar Plus plate reader (BMG Labtech). Enzymatic activity was calculated using the change in absorbance over time, where the concentration = Δabsorbance at 405 nm/(path length x extinction coefficient). The extinction coefficient used was 18000 M^-1^cm^-1^, and path length was 0.57 cm.

### Quantifying phosphate concentrations in the medium and calculating phosphate uptake rates

For phosphate detection in the media, a BIOMOL Green kit (Enzo Life Sciences, Germany) was used following the manufacturers protocol. 1:1 serial dilutions of the phosphate standard in ddH_2_O from the kit (800 μM) were made. The assay buffer was filtered seawater. Another standard using the phosphate from f/2 (36 μM) was used in a 1:1 serial dilution for comparison. Experimental samples were added to a 96-well plate and 100 μl BIOMOL^®^ GREEN reagent added. The samples were left to incubate at room temperature for 30 mins and read on a spectrophotometer plate reader at OD_620_ nm. Phosphate uptake rates were measured as previously described by (Harrison et al., 2025). Briefly, a medium sample was taken to confirm the initial level of phosphate, 500 µl of sample was removed and centrifuged (10,000 *g*, 5 min) and 200 µl of supernatant retained for analysis. Then phosphate was resupplied to cultures including WT and mutant lines, and 500 µl of sample centrifuged (10,000 *g*, 5 min) and 200 µl of supernatant retained for analysis at increments between 0-90 mins. The rate of phosphate uptake was calculated by subtracting the phosphate detected in the media from 36 µM at the different time points, and calculating the change in phosphate in the cell fraction per min. This was then normalised to the cell density to get pM per cell min^-1^.

## Results

### *In silico* analysis of *P. tricornutum* CDPK repertoire, structure and phylogeny

CDPKs have been identified previously in *P. tricornutum* transcriptomic datasets, with several genes showing increased transcriptional expression in P-deplete versus replete conditions (Alipanah et al. 2015; Cruz de Carvalho et al. 2016; Sharma et al. 2020). However, full analysis of *P. tricornutum* CDPKs, their structure and phylogeny, has not been reported. We therefore performed a targeted search for CDPKs in the *P. tricornutum* genome. Sequence similarity searches querying functionally characterised *A. thaliana* AtCPK1 (**Methods**), yielded nine CDPK hits containing both the necessary kinase and EF hand domain/s (**Figure 1A**). Five of these hits had four predicted EF-hands typical of canonical CDPKs (Bredow and Monaghan 2019). One of which (protein id. 12820) was annotated previously in the genome as PtCDPK1. The remaining four hits also had kinase domains, but two or fewer predicted EF-hands. An unrooted maximum-likelihood phylogenetic tree was reconstructed to determine the phylogenetic relationship of *P. tricornutum* CDPK-like sequences to homologs in other photosynthetic eukaryotes (diatoms, plants and green algae). This analysis revealed three distinct clades in diatoms (I-III), with at least one *P. tricornutum* CDPK hit in each clade. Clade II includes all five canonical CDPKs as well as Pt23694, none of which grouped with plant-specific CDPKs or CDPK-related kinases CRK clades (**Figure 1B**) (Bredow and Monaghan 2022). Pt25067 and Pt1875, which each have two predicted EF-hands, belong to distinct diatom clades III (Pt25067) and I (Pt1875). To investigate the potential role of the different *P. tricornutum* CDPKs during P limitation, we examined their expression in a previously published transcriptomic study (Cruz de Carvalho et al. 2016). Gene expression data in this analysis was available for all nine of the *P. tricornutum* CDPK hits. Three of these genes *PtCDPK2*, *PtCDPK3* and *PtCDPK5* showed a significant (p<0.05) greater than two-fold increase in expression after 4- and 8-days of P starvation relative to four-day old replete control treatment (**Figure 1C**) (Cruz de Carvalho et al. 2016). All three have multiple PtPSR1 recognition motifs in their promoter regions (Sharma et al. 2020). Amongst these, *PtCDPK2* showed the strongest transcriptional upregulation under P depletion, and its gene expression pattern strongly resembled that of *PtPSR1*.

### PtCDPK2 shows plasma membrane localisation and is tightly co-regulated with PtPSR1

Given the strong transcriptional upregulation of *PtCDPK2* by P depletion, we further characterised PtCDPK2 by determining its sub-cellular localisation, and examining its regulation by P availability, including in relation to PtPSR1. We generated transgenic *P. tricornutum* expressing PtCDPK2 tagged at the C-terminus with the fluorescent protein mVenus under the control of its upstream promotor region (**Methods**). A PtPSR1-mVenus line (Harrison et al. 2025) was also examined for comparison. Transgenic lines expressing PtCDPK2-mVenus and PtPSR1-mVenus were grown in P replete (36 µM initial phosphate, as in f/2 medium) and previously defined P-limiting (1.8 µM initial phosphate) conditions (Helliwell et al. 2021a). PtCDPK2-mVenus was strongly expressed under P limitation, but no fluorescence was detectable in P-replete conditions (**Figure 2A**). PtCDPK2-mVenus localised primarily to cell periphery, suggesting an association with the plasma membrane. This was confirmed by co-staining with the membrane dye FM4-64 (**Figure S1**). PtCDPK2 does not have a transmembrane domain, nor an N-terminal glycine for myristylation. Myristylation is a key post-translational modification involved in membrane targeting of many plant CDPKs, (Boudsocq and Sheen 2013; Yip Delormel and Boudsocq 2019), albeit several membrane-bound CPKs do lack this motif indicating the importance of other targeting mechanisms too (Li et al. 2008; Zhang et al. 2014; Wen et al. 2020). In addition to PtCDPK2-mVenus, we examined PtPSR1-mVenus cells. In concurrence with transcriptional expression (Sharma et al. 2020), PtPSR1-mVenus was strongly upregulated under P limitation and localised to the nucleus (**Figure 2B**), as previously reported (Harrison et al. 2025).

**Figure 2.**
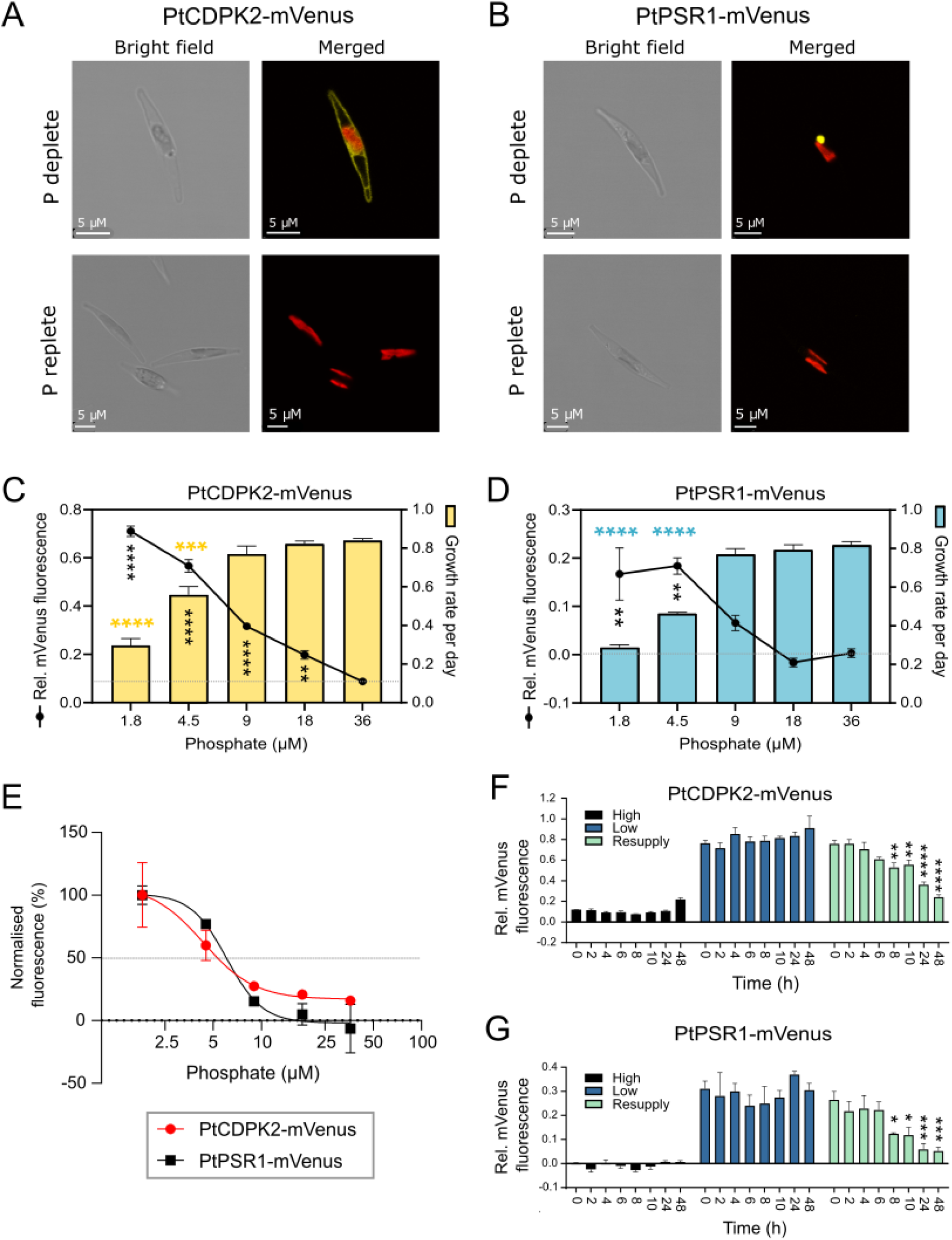
PtCDPK2 is plasma membrane localised and tightly co-regulated with PtPSR1 by P availability. **A)** Under P deplete conditions (f/2 medium with 1.8 µM phosphate) PtCDPK2-mVenus is localised to the plasma membrane with no fluorescence detected in P replete conditions (36 µM Pi) on day 4. **B)** Expression and localisation of PtPSR1-mVenus in P deplete versus replete conditions (day 4). **C** and **D)** Growth rate (day^-1^) between day 4 and 5 and relative mVenus fluorescence (mVenus/chlorophyll fluorescence) on day 5 for the PtCDPK2-mVenus (**C**) and PtPSR1-mVenus (**D**) strains. Data are mean ± standard error. Statistically significant results of a one-way ANOVA analyses comparing growth rate (day^-1^) of each phosphate treatment to the 36 µM control via a Dunnett’s multiple comparisons test, are indicated with coloured horizontal asterisks (**** p<0.0001; *** p<0.001; ** p<0.01; * p<0.05). The same test was also performed on the relative mVenus fluorescence data, and the results are shown with black vertical asterisks. **E)** Fluorescence of PtCDPK2-mVenus and PtPSR1-mVenus at different phosphate concentrations on day 4 as a percentage of that measured at 1.8 µM phosphate. Sigmoidal dose response curves were fitted using a 4-parameter log-logistic model (Mehrshahi et al., 2020). Data are presented as mean ± standard deviation (n=3). **F** and **G)** Time-course of relative mVenus fluorescence (mVenus/chlorophyll fluorescence) of PtCDPK2-mVenus (**F**) and PtPSR1-mVenus (**G**) strains grown under P replete (36 µM Pi), deplete (1.8 µM Pi) and following resupply of 36 µM phosphate to 4-day old deplete cells. Results of a two-way ANOVA are indicated, comparing relative mVenus fluorescence at each time-point to that at time 0 (i.e. prior to P resupply to the resupply cultures), for each strain within the different P treatments (**** p<0.0001; *** p<0.001; ** p<0.01; * p<0.05).

We next measured the expression of PtCDPK2-mVenus and PtPSR1-mVenus over time when inoculated into media with different starting concentrations of phosphate. Increases in relative PtCDPK2-mVenus and PtPSR1-mVenus fluorescence (mVenus fluorescence normalised to chlorophyll fluorescence) were clearly visible by day 4 in treatments with initial phosphate concentrations of 1.8 µM and 4.5 µM (**Figure S2**). For higher phosphate conditions, including 9 µM and 18 µM, there was no increase in relative mVenus fluorescence at day 4, but increases were seen at later timepoints. Even with 36 µM phosphate, some mVenus expression was detectable by day 7 for PtCDPK2-mVenus. In contrast, relative fluorescence of control WT cells remained around the baseline (of zero) for all phosphate concentrations throughout the experiment (**Figure S2**). Examining growth rate (day^-1^) versus mVenus fluorescence for the two strains across the range of phosphate concentrations tested, demonstrated that PtCDPK2-mVenus and PtPSR1-mVenus fluorescence was most strong when growth was P limited (**Figure 2C-D**). However, for PtCDPK2-mVenus, significant increases in relative fluorescence (black asterisks; p-value<0.01, one-way ANOVA) were also evident even when there were no significant declines in growth rate compared to the 36 µM phosphate condition (**Figure 2C**). Together, these results demonstrate that P starvation alone i.e. prior to growth limitation by P, can trigger upregulation of PtCDPK2. Moreover, PtCDPK2-mVenus and PtPSR1-mVenus show very similar expression dynamics. Plotting mVenus fluorescence of PtCDPK2-mVenus versus PtPSR1-mVenus for each individual phosphate concentration on day 4 as a percentage of that in the 1.8 µM phosphate treatments further demonstrated this (**Figure 2E**), with the level of initial phosphate in the medium needed for 50% expression calculated as 4.3 and 6.1 µM phosphate, respectively.

Having demonstrated strong increases in PtCDPK2-mVenus and PtPSR1-mVenus under P limitation, we monitored fluorescence following phosphate resupply. Significant reductions by 30% and 54% for PtCDPK2-mVenus and PtPSR1-mVenus respectively, were observed within 8 h of phosphate resupply to 4-day old 1.8 µM grown cells (P<0.05, two-way ANOVA) (**Figure 2F-G**). Values were comparable to replete cells within 48 h. To investigate the Ca^2+^ dependency of this response. We applied a pharmacological inhibitor of the P-Ca^2+^ signalling response, RuR (5 µM) (Helliwell et al. 2021a), to P-deplete cells prior to phosphate resupply on day 4 (**Figure S3A-B**). Replete and deplete cells were also examined in the presence and absence of RuR as controls. Decreases in PtCDPK2-mVenus fluorescence were observed after 6 h and 24 h following phosphate resupply in -RuR and +RuR pretreated cells in a similar manner (**Figure S3C-D**). This indicates the regulation of PtCDPK2 following phosphate resupply to P-limited cells is not dependent on P-Ca^2+^ signalling.

### A threshold intracellular P level defines activation of P-Ca^2+^ signalling

Phosphate-induced Ca^2+^ signals are also apparent only in P-deplete and not P replete cells (Helliwell et al. 2021a). As PtCDPK2 may play a role in sensing Ca^2+^ elevations generated by phosphate resupply, we examined whether P-Ca^2+^ signalling is activated by P-starvation in a similar manner to PtCDPK2. We grew PtR1 cells expressing the Ca^2+^ indicator R-GECO1 in f/2 medium with different initial phosphate concentrations, including: 1.8, 4.5, 9, 14, 18, 27 or 36 µM. The capacity of cells for phosphate induced Ca^2+^ signalling was then examined via Ca^2+^ imaging on day 4 and 5. After 4 days, no cells in the 36 µM phosphate treatment responded to phosphate resupply, with only a small percentage responding at the slightly lower concentration of 27 µM (**Figure 3A**). A significant increase in the normalised mean maximal fluorescence intensity (F/F_0_) in response to phosphate resupply compared to the 36 µM replete control was detectable for all treatments with 18 µM phosphate or lower (Kruskal-Wallis test, p-values <0.001). By day 5, there was a significant increase (p-value <0.01) in normalised mean maximal F/F_0_ even in 27 µM phosphate grown cells (**Figure 3B**). Notably, exogenous phosphate was undetectable in all treatments by day 4, including the 36 µM control. This indicates that exhaustion of phosphate from the external environment does not trigger activation of the capacity of cells for P-Ca^2+^ signalling. As no external phosphate was detected, we assumed all phosphate had been taken up by the cells, allowing calculation of cellular phosphate quotas by dividing the initial phosphate concentration by cell number. Phosphate quotas per cell declined with decreasing levels of phosphate in the initial medium, as would be expected (**Figure 3C-D**). Cellular phosphate quotas became further depleted by day 5 compared to day 4 in all treatments. Notably, cellular phosphate needed to deplete to around 0.5 pg phosphate/cell for cells to exhibit P-Ca^2+^ signalling on both day 4 and 5. Together, this demonstrates that a threshold intracellular P level defines activation of the capacity for P-Ca^2+^ signalling and likely other P-signalling responses. Moreover, activation of the ability for P-Ca^2+^ signalling occurred before limitation of growth rate by P, in a similar manner to increases in PtCDPK2-mVenus expression (**Figure 3D-E**).

**Figure 3.**
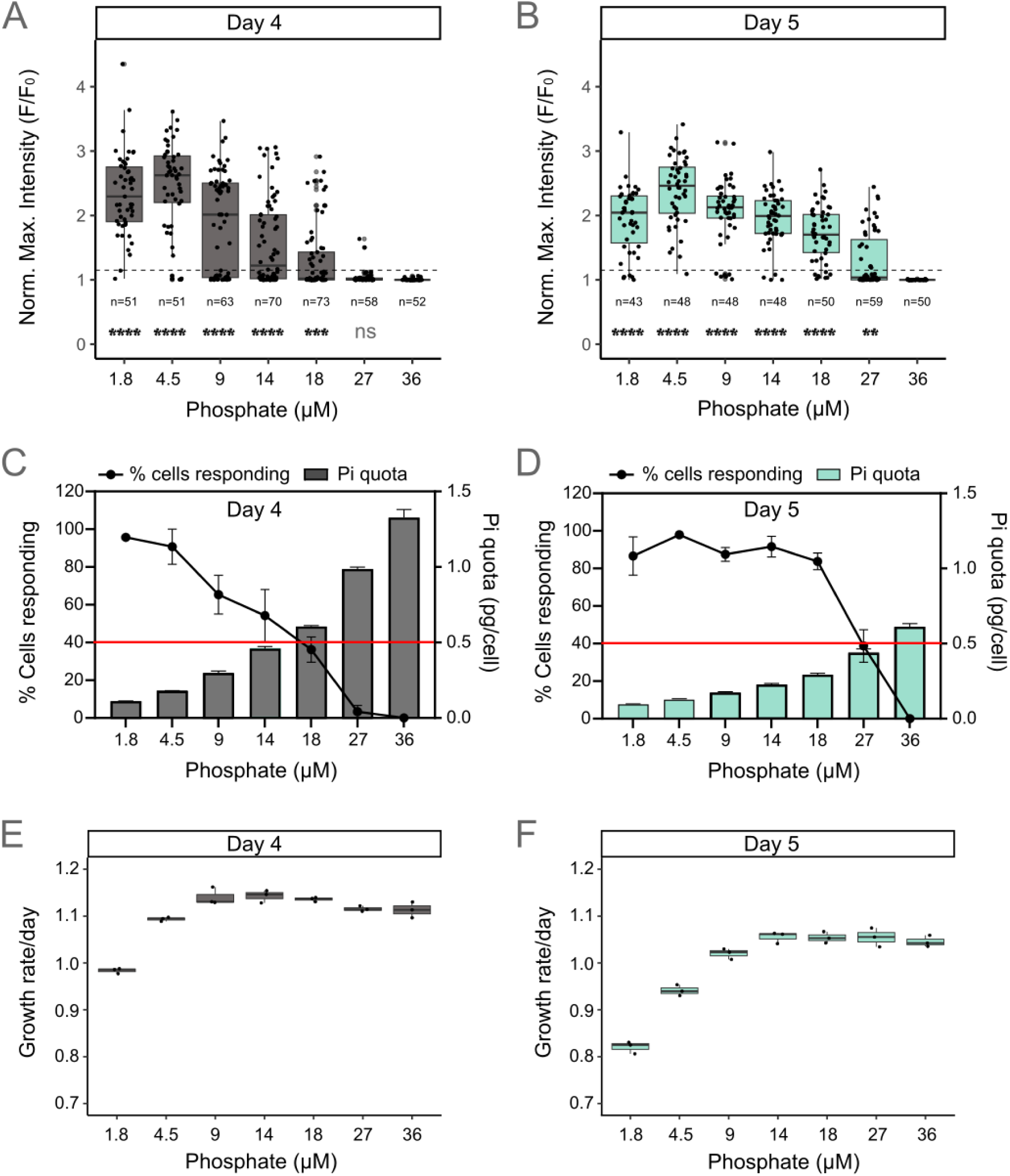
A threshold intracellular P level defines activation of P-Ca^2+^ signalling. **A**) Mean normalised maximal fluorescence (F/F_0_) of PtR1 cells grown in different starting concentrations of phosphate and resupplied with 36 µM phosphate and day 4, and day 5 (**B**). Data is compiled from three independent replicate experiments per treatment, and the total no. of cells examined per treatment is indicated on the figure. Results of a non-parametric Kruskal-Wallis test comparing each treatment to the P replete 36 µM control, are also given: *** p<0.0001; *** p<0.001; ** p<0.01; * p<0.05 and ‘ns’ not significant. **C**) and **D**) Phosphate quota per cell (pg per cell) after 4 and 5-days respectively (calculated by dividing the initial phosphate concentration by cell number). The percentage of cells showing a phosphate induced Ca^2+^ signalling response is also shown (black line). Cells were deemed to have responded if the [Ca^2+^]_cyt_ fluorescence signal was above the F/F_0_ threshold of 1.15. The red line indicates where 40% of cells respond. Data are presented as mean ± standard error (n=3). **D**) and **E**) Growth rate (day^-1^) between day 0 and 4 (D) and 0 and 5 (E) of the PtR1 strain in the different phosphate treatments. Data are presented as mean ± standard error (n=3)

### Generation of biallelic *Ptcdpk2* mutants

To further investigate the functional role of *PtCDPK2*, mutant lines were generated using CRISPR-Cas9 gene editing. Two single guide RNAs (**Figure S4A**, Methods) were used, targeting the region of the gene encoding the protein kinase domain. Putative mutants were screened via PCR band-shift assays with primers flanking the sgRNA target sites (**Figure S4A-B**). Four independent biallelic mutants were obtained showing single bands either larger (*Ptcdpk2.1*), or smaller (*Ptcdpk2.3*, *Ptcdpk2.4*, and *Ptcdpk2.5*) compared to WT. Analysis via Sanger sequencing showed *Ptcdpk2.1* possesses an insertion of 109 bp (**Figure S4A**). *Ptcdpk2.3* and *Ptcdpk2.4* have single deletions of 318 bp and 546 bp respectively, whereas *Ptcdpk2.5* has two separate deletions 284 bp and 43 bp in size. A single nucleotide polymorphism (SNP) was additionally identified in *Ptcdpk2.3.* To determine the impacts of these modifications on the coding sequence we generated cDNA via RT-PCR analysis (**Figure S4C**). Sanger sequencing of cDNA indicated that the insertion in the coding sequence of *Ptcdpk2.1* causes a frameshift and in-frame stop codon, severely truncating the protein (**Figure S4D**). In contrast, *Ptcdpk2.3* and *Ptcdpk2.5* each had distinct deletions within the region of the transcript encoding the kinase domain. The SNP in *Ptcdpk2.3* would cause an alanine (Ala) to threonine (Thr) substitution in a conserved region of the protein. Finally, RT-PCR of *Ptcdpk2.4* revealed multiple transcripts, suggesting alternative splicing, although all variants analysed (**Supplementary Methods**) were predicted to produce severely truncated proteins (**Figure S4C-D**).

### *Ptcdpk2* mutants show increased physiological stress under phosphate limitation

*Ptcdpk2* mutants were examined under P replete and deplete conditions (36 µM and 1.8 µM of initial phosphate, respectively) and following phosphate resupply. For the phosphate resupply experiment 1.8 µM phosphate grown cells were resupplied with 36 µM on day 4 and growth monitored for all treatments between day 4 and 5. Growth dynamics of *Ptcdpk2* mutants and WT were comparable between P regimes, with no statistically significant differences detected in the growth rate of mutants in low or high phosphate regimes (**Figure 4A**). Following phosphate resupply, growth rate did not differ between the WT and mutants either, suggesting similar growth recovery capacity across all strains. Moreover, all mutants show significant recovery of photosynthetic efficiency (F_v_/F_m_) values within 24 h of phosphate resupply, like WT (p-value< 0.0001; two-way ANOVA, **Figure 4B**). However, when resupplying phosphate on day 6 instead of day 4 (and monitoring growth rate between day 6 and 7), we noticed that all mutants showed impaired growth rate recovery following phosphate resupply compared to WT (**Figure 4C**).

**Figure 4.**
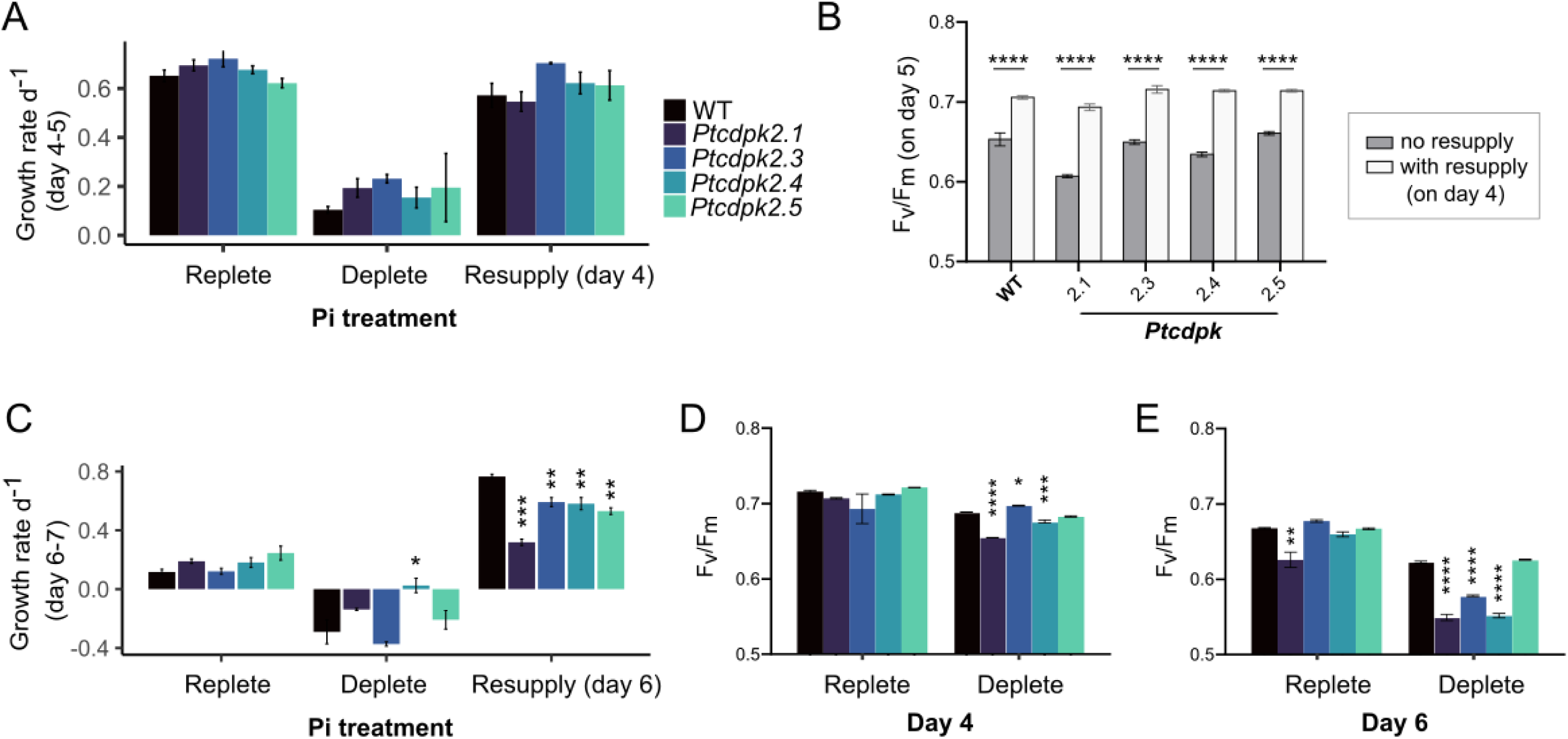
*Ptcdpk2* mutants show increased physiological stress under phosphate (Pi) limitation. **A)** Growth rate (day^-1^) between day 4 and 5 for samples grown under different Pi regimes, including: replete (36 µM), deplete (1.8 µM) and following resupply of 36 µM Pi to 4-day old deplete cells. Data are presented as mean ± standard error (n=3). A two-way ANOVA was performed on growth rates of mutants compared to WT within each different treatment, but the data was not significant. **B)** F_v_/F_m_ values of cultures on day 5 with or without resupply on day 4. Data are presented as mean ± standard deviation (n=3). Results of a two-way ANOVA comparing the resupply versus no resupply treatment for each line: * p-value <0.05, ** p-value <0.01, *** p-value <0.001, **** p-value <0.0001. **C)** As in (A) but growth rate (d^-1^) between day 6 and 7, with Pi resupplied on day 6 instead of day 4. Data are presented as mean ± standard error (n=3). Results of a one-way ANOVA and post-hoc Tukey test are shown, comparing growth of mutants versus WT within each different treatment: * p-value <0.05, ** p-value <0.01, *** p-value <0.001. **D)** Photosynthetic efficiency (F_v_/F_m_) of WT compared to mutants in cultures inoculated into replete (36 µM phosphate) or deplete (1.8 µM phosphate) conditions on day 4. Data are presented as mean ± standard error (n=3). Results of a two-way ANOVA comparing each *Ptcdpk2* mutant to WT within each treatment: * p-value <0.05, ** p-value <0.01, *** p-value <0.001, **** p-value <0.0001. **E**) As in D, but on day 6.

To better understand this, we further examined F_v_/F_m_ values of mutants versus WT in P replete and deplete conditions, on day 4 and 6. No statistically significant differences were observed between WT and *Ptcdpk2* mutants grown in replete conditions on day 4 (**Figure 4D**; two-way ANOVA). However, F_v_/F_m_ values of deplete cultures were significantly lower in mutants *Ptcdpk2.1*, *Ptcdpk2.3 and Ptcdpk2.4* compared to WT, and this effect became more pronounced by day 6 (**Figure 4E**). Whilst there was also a significant reduction in F_v_/F_m_ of *Ptcdpk2.1* compared to WT on day 6 for 36 µM grown cells, data from the previous experiment indicates that 36 µM grown cells at day 5 are near the threshold of intracellular P depletion for induction of P-Ca^2+^ signalling (**Figure 3D**) and are likely experiencing P starvation by day 6. Together these experiments demonstrate that *Ptcdpk2* mutants are defective in their ability to cope with P limitation, rather than the phosphate resupply recovery response. When experiencing more severe P depletion (i.e. after 6 days in low P medium) cells are less able to recover following phosphate resupply, but this is because mutants are experiencing enhanced physiological stress prior to resupply compared to WT.

### *Ptcdpk2* mutants retain Ca^2+^ signalling in response to phosphate resupply

We have demonstrated that PtCDPK2 is plasma membrane localised and shows similar induction of expression in response to P depletion compared to the activation of the P-Ca^2+^ signalling response (**Figures 2** and **3**). In plants, membrane associated CDPKs can play crucial roles in ion channel activation (Mori et al. 2006; Mehlmer et al. 2010; Brandt et al. 2012). We have previously demonstrated that phosphate resupply induced Ca^2+^ signals are dependent on external Ca^2+^ levels, suggesting that plasma membrane channel/s are involved (Helliwell et al. 2021a). We therefore examined the ability of *Ptcdpk2* mutants for P-Ca^2+^ signalling. We transformed *Ptcdpk2* mutants with the ratiometric fluorescent biosensor R-GECO1-mTurquoise (RGMT) (**Methods**) (Harrison et al. 2025), and obtained fluorescent strains for *Ptcdpk2.1, Ptcdpk2.3* and *Ptcdpk2.4*. WT cells expressing RGMT were used as a control. As expected, P-replete WT RGMT cells did not respond to phosphate resupply (**Figure 5A**, left). The same was also the case for the three mutants examined (**Figure 5B-D**). In contrast, phosphate resupply to P-deplete cells yielded a rapid elevation in cytosolic Ca^2+^ in WT RGMT (**Figure 5A**, right). This was also the case for all the mutants, although the percentage of cells responding above a threshold R/R_0_ of 1.15 was lower for *Ptcdpk2.1*. The maximal intensity of those *Ptcdpk2.1* cells that did respond, was significantly reduced compared to WT unlike the other *Ptcdpk2* lines. However, together the results suggest that PtCDPK2 is not essential for P-Ca^2+^ signalling. This is in keeping with the ability of *Ptcdpk2* mutants for apparently normal physiological recovery following phosphate resupply on day 4 (**Figure 4**).

**Figure 5.**
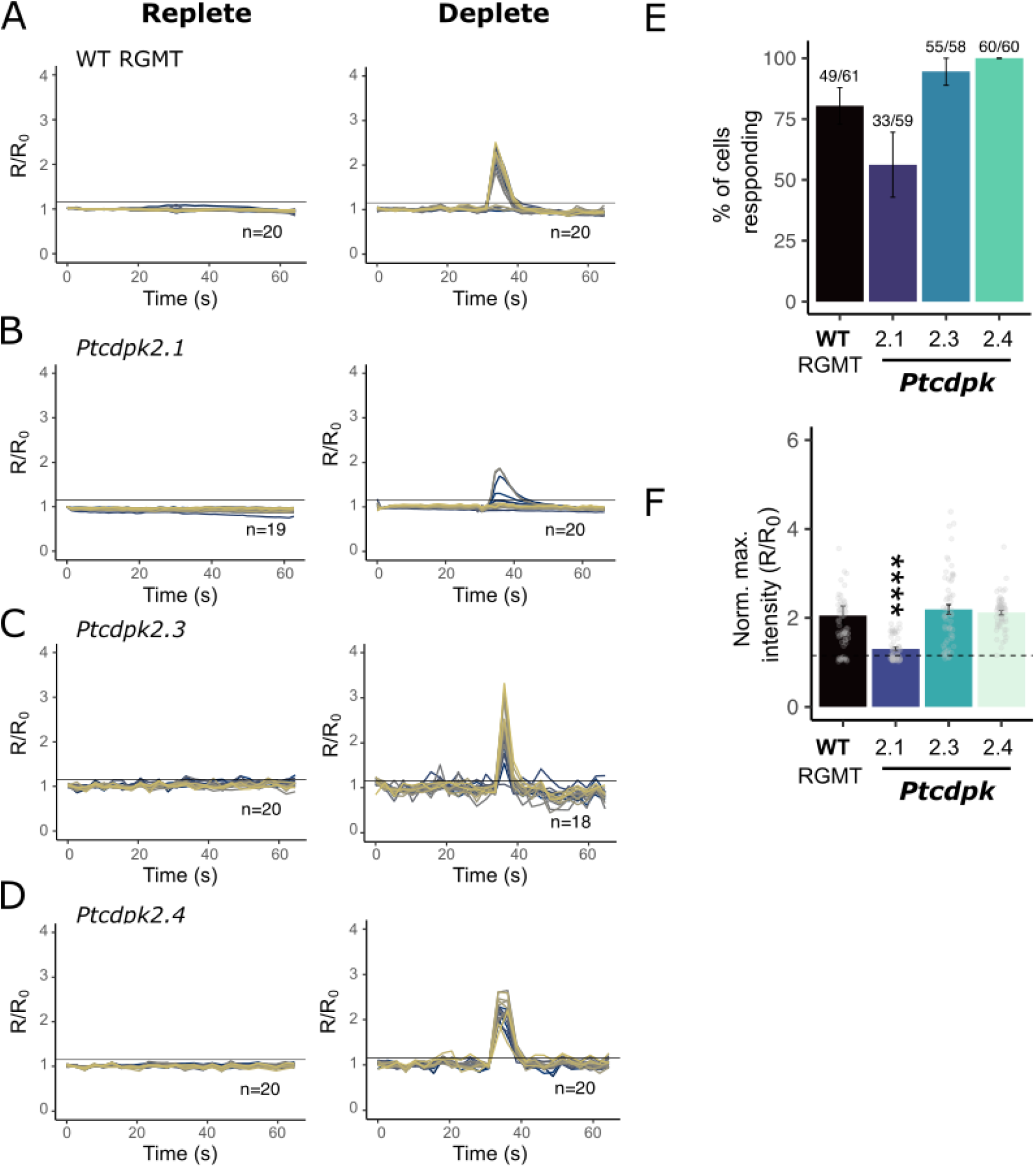
Phosphate-induced Ca^2+^ signalling in *Ptcdpk2* mutants compared to wildtype. R-GECO1-mTurquoise (RGMT) was expressed in *Ptcdpk2* mutants, as well as wildtype (WT). Cells were grown in high (36 µM) or low phosphate (1.8 µM) for 4 days. Live-cell imaging experiments of cells perfused with f/2 medium with no phosphate for 30 s and then with f/2 medium with 36 μM phosphate for 30 s were conducted. Representative Ca^2+^ signalling trace graphs showing R/R_0_ for high and low Pi grown four-day old cells **A)** WT (*RGMT*), **B)** *Ptcdpk2.1* **C)** *Pt*c*dpk2.3* and **D)** *Ptcdpk2.4*. R_0_ is the R value at the start of the experiment (I.e. time 0) and R is the ratio of R-GECO1/mTurquoise fluorescence. **E)** Percentage of cells responding in WT and mutant lines grown in P deplete conditions, with a normalised max fluorescence intensity above the threshold of 1.15. Data compiled from three independent replicate treatments per line. No. of cells responding/total no. of cells examined is given. **F)** Normalised max intensity (R/R_0_) of cells that did respond in the P deplete treatment. Data compiled from three independent trace graph experiments (in A). Statistically significant results of a non-parametric Kruskal-Wallis one-way ANOVA test comparing mutant lines to WT RGMT are indicated: **** p-value <0.0001.

### *Ptcdpk2* mutants have aberrant alkaline phosphatase activity and phosphate uptake capacity under P starvation

To further investigate what might be causing the reduced cell health of mutants compared to WT, we examined P scavenging (alkaline phosphatase activity) and uptake capacity in the different strains. All four *Ptcdpk2* mutants showed reduced alkaline phosphatase activity on day 3 compared to the WT, and this was statistically significant for *Ptcdpk2.1*, *Ptcdpk2.3* and *Ptcdpk2.4* (p-value <0.001; one-way ANOVA, **Figure 6A**). Whilst the effect was less pronounced by day 4, enzyme activity remained significantly lower for *Ptcdpk2.1* and *Ptcdpk2.3*. As no significant differences in growth of mutants versus WT were observed in these conditions (**Figure 4A**), the changes in enzyme activity must be driven by the ability of mutants for alkaline phosphatase activity under P depletion, rather than due to differences in growth stage.

**Figure 6.**
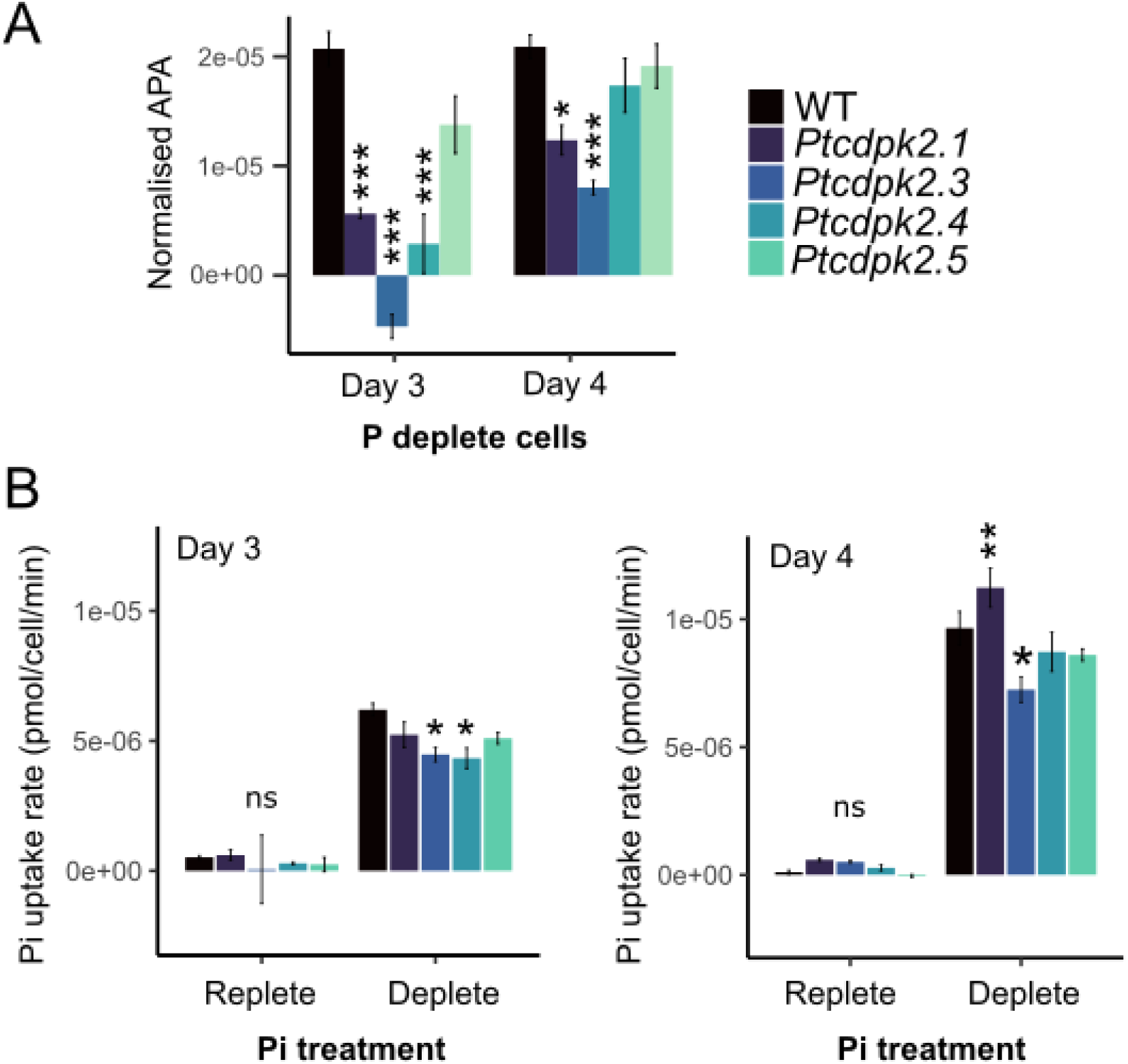
*Ptcdpk2* mutants show aberrant alkaline phosphatase activity and phosphate (Pi) uptake capacity during P starvation. **A)** Alkaline phosphatase activity (APA) (pNPP fmol/min) normalised to chlorophyll fluorescence of cultures grown in f/2 medium with low phosphate (1.8 µM) after 3 and 4 days. Data are presented as mean ± standard error (n=3). Data were tested for normality using the Shapiro test and a one-way ANOVA was performed comparing mutants to WT with Tukey post-hoc test: * p-value <0.05, *** p-value <0.001. **B)** Pi uptake rates (pmol/cell/min) of 3-day (left) and 4-day (right) old cells grown in P replete (36 µM phosphate) and P deplete (1.8 µM phosphate) conditions. Results of a one-way ANOVA comparing mutants to the WT control are shown: * p-value <0.05, ** p-value <0.01). Data are presented as mean ± standard error (n=3).

We also quantified phosphate uptake capacity of the different strains pre-grown in medium with either 36 µM or 1.8 µM phosphate. Cells were then resupplied with 36 µM and depletion of phosphate from the medium measured to ascertain phosphate uptake rates. Whereas no differences were seen in the uptake capacity of strains grown in P replete conditions, significant differences were detected between WT and *Ptcdpk2.3* and *Ptcdpk2.4* on day 3, as well as *Ptcdpk2.1,* and *Ptcdpk2.3* on day 4 of P limitation (p-value <0.05; one-way ANOVA, **Figure 6B**). Hence, our results show that three of the four *Ptcdpk2* mutants have evidence of aberrant alkaline phosphatase activity and phosphate uptake capacity. These phenotypes were only observed in P deplete conditions and therefore likely account for the reduced cell health of mutants in low P medium compared to WT. The apparently unimpaired alkaline phosphatase activity and phosphate uptake rates of *Ptcdpk2.5* likely explains why this mutant does not exhibit significant declines in F_v_/F_m_ compared to WT under P depletion (**Figure 4D**).

## Discussion

CDPKs have been studied in divergent eukaryotes including plants, green and red algae (Valmonte et al.; Brawley et al. 2017), and heterotrophic protists (Billker et al. 2009; Sharma et al. 2021). Here, we report a comprehensive examination of CDPKs in the diatom *P. tricornutum*. We identified five canonical *CDPK* genes, encoding both the expected kinase domain as well as four EF hands (Harper et al. 2004; Bredow and Monaghan 2022). Diatom CDPKs were phylogenetically distinct from plant and green algal CDPKs (Li et al. 2019). Three *P. tricornutum CDPK*s are strongly upregulated by P limitation (Cruz de Carvalho et al. 2016; Sharma et al. 2020; Villar et al. 2025), and all contain PtPSR1 recognition motifs in their promoter regions (Sharma et al. 2020). We focussed on characterising *PtCDPK2*, which showed the greatest transcriptional upregulation by P starvation. We hypothesised that PtCDPK2 may be involved in sensing P-induced Ca^2+^ elevations and coordinating recovery responses of P-limited *P. tricornutum* cells. However, our collective evidence instead supports a role for PtCDPK2 in the regulation of P-starvation physiology, with *Ptcdpk2* mutants showing significantly reduced F_v_/F_m_, alkaline phosphatase activity and aberrant phosphate uptake under P depletion. PtCDPK2 protein abundance is strongly induced by P limitation, and rapidly downregulated following phosphate resupply. This expression profile strongly resembles that of PtPSR1, which likely regulates *PtCDPK2* (Sharma et al. 2020). Hence, we propose PtCDPK2 confers a second tier, or outpost of PtPSR1, in regulating *P. tricornutum* P-starvation responses. Whilst exhibiting similar expression dynamics, these proteins are spatially separated. PtCDPK2 is detected in the vicinity of the plasma membrane, whereas PtPSR1 is nuclear localised (Harrison et al. 2025). Thus, PtCDPK2 likely affords regulatory roles that the nuclear localised PtPSR1 transcription factor cannot fulfil, such as controlling localisation, activity and/or interactions with other proteins on or near the plasma membrane. As we found no evidence that PtCDPK2 is involved governing downstream P resupply recovery, the Ca^2+^ sensors/decoders in this pathway remain to be determined.

The identification of five canonical CDPKs and four CDPK-like genes in *P. tricornutum* is considerably fewer than reported for green algae and plants, with CDPK abundance increasing with morphological complexity in these lineages (Edel et al. 2017). *C. reinhardtii* encodes 15 CDPKS, whereas *A. thaliana* has 34 (Yip Delormel and Boudsocq 2019). These CDPKs have broad functional roles including in nutrient transport and signalling, but also flagella biogenesis, plant growth and development, immune signalling, alongside tolerance regulation to abiotic and biotic pressures (Liang and Pan 2013; Motiwalla et al. 2014; Liu et al. 2017; Shi et al. 2018; Yip Delormel and Boudsocq 2019; Qin et al. 2020). A major challenge limiting study of plant CDPKs has been functional redundancy (Liu et al., 2017; Yip Delormel yip & Boudsocq, 2019b). Certainly, the other *P. tricornutum* CDPKs upregulated under P limitation could perform redundant roles in *Ptcdpk2* mutants. Clearly complete compensation did not occur, since we observed convincing reductions in the F_v_/F_m_ of mutants compared to WT under P limitation. Notably, the CDPK repertoire of *P. tricornutum* is more comparable in size to apicomplexan protists, which are more closely related to diatoms than plants and green algae (Keeling and Burki, 2019b). *Plasmodium falciparum* has seven, and *Toxoplasma gondii* around twelve CDPKs, three of which are CDPK-like with 3 EF-hands akin to plant CCAMKs (Patil et al. 1995; Billker et al. 2009). Apicomplexan CDPKs can be essential for parasite viability, necessitating the use of conditional knockout and/or chemical inhibition to facilitate functional gene characterisation (Kato et al. 2008; Lourido et al. 2010). Integrative techniques combining protein localisation, reverse genetics, expression analysis and phosphoproteomics (Liu et al. 2020) will help illuminate further the cellular roles of other diatom CDPKs. However, there is a clear need for molecular tool development enabling conditional knockout to allow generation of double and triple mutants and help overcome the potential for gene lethality/redundancy in the *P. tricornutum* system.

The localisation of PtCDPK2 to the cell periphery, strongly suggests a primary role regulating membrane processes. *P. tricornutum* encodes eight alkaline phosphatases and ten phosphate transporters (Dell’Aquila et al. 2020). Alkaline phosphatases PtPhos5, PtPhos6 and PtPhos7 are endomembrane localised, whereas PtPhos8 and PtPhos3 are in the plasma membrane and endomembranes (Dell’Aquila et al. 2020). Notably, PtPhos1 and PtPhos2 are secreted (Lin et al. 2013, 2017; Buhmann et al. 2016; Dell’Aquila et al. 2020). Whilst PtPhos1 is detected in P replete and deplete conditions it is secreted only under low P. These dynamics are hypothesised to be modulated by protein phosphorylation (Dell’Aquila et al. 2020), but the specific phosphosites and kinase/s responsible are yet to be deduced. Perturbed alkaline phosphatase activity in *Ptcdpk2* mutants points to PtCDPK2 as a strong candidate. *P. tricornutum* phosphate transporters are also distributed in different cellular locations, with Na^+^/phosphate symporters PtNap2, PtNap4 and PtNap5 plasma membrane based, like PtCDPK2. Additionally, PtNap3 and the H^+^/Pi transporter PtHP1 are present in the plasma membrane and distinct spot-like structures (Dell’Aquila et al. 2020). A vacuolar phosphate transporter (PtVPT1) has also been reported. Protein phosphorylation is crucial for trafficking phosphate transporters within plant cells. The high affinity transporter PHT1 requires phosphorylation at Ser-514 for export from the ER to the plasma membrane via interactions with PHOSPHATE TRANSPORTER TRAFFIC FACILITATOR1 (PHF1) (Bayle et al. 2011). Whilst

PtCDPK2 likely controls the activity and/or localisation of P transport/scavenging machinery or their interacting proteins, the significant redundancy in diatom P acquisition machinery introduces potential for phenotype compensation. Certainly, functional complementation of the secreted alkaline phosphatase (PtPhos1) by an intracellular enzyme (PtPhos6), and vice versa, has been reported previously in *P. tricornutum* (Zhang et al. 2022). This could explain why the phenotypes relating to alkaline phosphatase became less pronounced with time in P deplete conditions.

The ability to cope with fluctuating P supply and regulate P starvation physiology accordingly, demands molecular mechanisms to sense P availability. In *Saccharomyces cerevisiae*, internal P levels are sensed via SPX (SYG1/Pho81/XPR1) domain containing proteins that directly bind inositol phosphates, triggering synthesis of poly-P (Lonetti et al. 2011; Wild et al. 2016; Gerasimaite et al. 2017). Diatoms have SPX proteins (Zhang et al. 2021), as well as Vacuolar Transport Chaperone homologues (PtVTC 1–4) (Schreiber et al. 2017), but only PtVTC2 has been localised to the vacuole (Dell’Aquila et al. 2020). However, diatoms employ PSR1, instead of the Pho80/Pho85/Pho4 regulon for P starvation regulation. Employing a reporter gene approach, we have been able to study with high temporal resolution the regulation of PtPSR1 and PtCDPK2 by P availability, in relation to P-Ca^2+^ signalling, and internal and external P levels. Our collective evidence indicates that growth need not be P limited, and a threshold cellular P level, rather than exhaustion of external P, defines activation of P-signalling. Efforts must now focus on coupling such dynamics in the context of cellular P reserves, such as poly-P. Lapointe et al. (2024) recently developed methodologies to visualise poly-P granules in vacuoles of the freshwater diatom, *Achnanthidium minutissimum* (Lapointe et al. 2024). Like *P. tricornutum*, *A. minutissimum* displays a remarkable capacity to take up phosphate from the external medium, driving rapid accumulation of poly-P. This so-called overplus response is common also to cyanobacteria (Li & Dittrich, 2019) and green algae (Plouviez et al. 2021; Zúñiga-Burgos et al. 2024). Poly-P levels persisted for at least two days following inoculation of *A. minutissimum* into -P medium.

Moreover, exponential growth was maintained despite P depletion for nearly 10 days (Lapointe et al., 2024). This is testament to the attributes of diatoms for coping with P deprivation. Central to the regulation of P-starvation responses in divergent photosynthetic eukaryotes, is PSR1 (Wykoff et al. 1999; Rubio et al. 2001; Sharma et al. 2020). Overexpression of CrPSR1 in *C. reinhardtii* induces luxury phosphate uptake (Slocombe et al. 2023). Moreover, in *P. tricornutum Ptpsr1* mutants exhibit impaired P-Ca^2+^ signalling (Harrison et al., 2025). This evidence combined with the tight co-regulation of PtCDPK2 with PtPSR1, and the presence of PtPSR1 binding motifs in upstream regions of PtCDPK genes (Sharma et al. 2020), demonstrates PtPSR1 likely regulates Ca^2+^ sensor machinery. Yet evidence so far suggests diatom-like P-Ca^2+^-signalling is not displayed by other photosynthetic eukaryotes (Pivato et al., 2024). Thus, hijacking of PSR1 for regulation of Ca^2+^-signalling apparatus has likely occurred in diatoms. This is not dissimilar to land plants, whereby plant mycorrhizal symbiosis genes have become wired to the PSR1 protein PHR2 in rice (Das and Gutjahr 2022; Das et al. 2022). These examples highlight P nutrition as a powerful driver shaping divergent biology and diverse strategies of photosynthetic eukaryotes to survive in environments with variable P, be they aquatic or terrestrial. As a central hub regulating such processes, PSR1 may well provide an exciting gateway for discovery of novel genes underlying such biology.

## Acknowledgements

This work was supported by Natural Environment Research Council (NERC) Independent Research Fellowship NE/R015449/2 and Biotechnology and Biological Sciences Research Council (BBSRC) New Investigator Grant BB/W006286/1. Yasmin Meeda is also grateful for a jointly funded PhD studentship between the University of Exeter and Marine Biological Association (Plymouth, UK).

## Author Contributions

YM, EH, KH, GW and AM planned and designed the research. YM, EH, KH, SW and AH performed experiments and analysed data. KH and YM interpreted data and wrote the manuscript. All authors approved the finalised manuscript.

