## Supplemental Figures for "A plasma membrane Ca^2+^-dependent protein kinase PtCDPK2 promotes phosphorus starvation resilience in *Phaeodactylum tricornutum*"

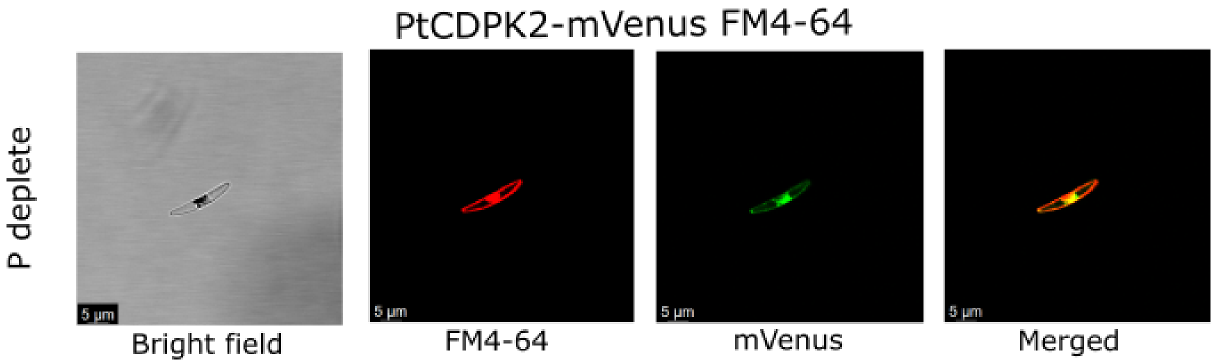


**Figure S1. PtCDPK2-mVenus expression and localisation under P-limiting conditions.** Localisation of PtCDPK2-mVenus and plasma membrane stain FM4-64 of 4-day old cells grown in low Pi (1.8 µM) medium and imaged using confocal microscopy.

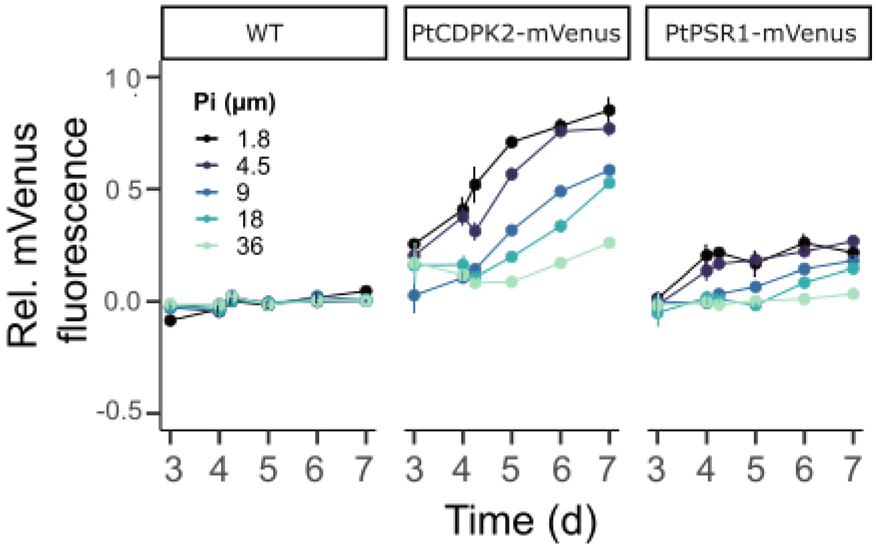


**Figure S2. PtCDPK2-mVenus and PtPSR1-mVenus are upregulated under P-starvation.** Relative mVenus fluorescence (mVenus/chlorophyll fluorescence) of wild-type (WT), PtCDPK2-mVenus and PtPSR1-mVenus strains over 7 days inoculated into different initial concentrations of phosphate (Pi). Data are presented as mean ± standard error (n=3).

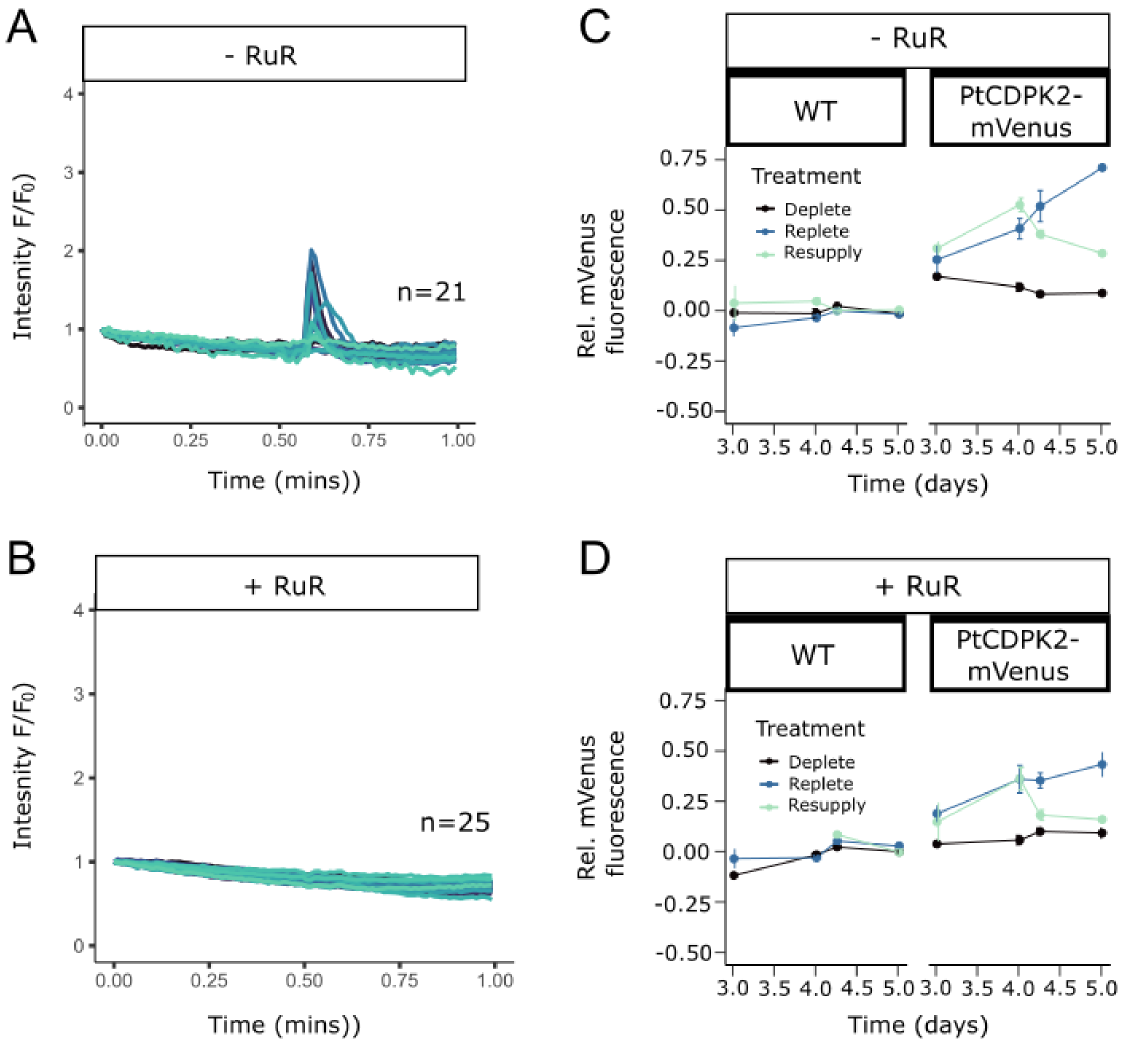


**Figure S3. Decreases in PtCDPK2-mVenus fluorescence following phosphate resupply to P deplete cells are not dependent on P-$Ca^{2+}$ signalling. A**) Fluorescence traces ($F/F_0$) of 4-day old R-GECO1 (PtR1) cells grown in P deplete (1.8 µM phosphate) conditions exposed to 36 µM phosphate resupply. Cells without RuR were perfused with filtered seawater (FSW) for 30 s and then FSW with 36 µM phosphate for 30 s. **B**) Experiment as described in (A) except cells were pre-treated with 5 µM ruthenium red (RuR) for 5 mins. **C**) PtCDPK2-mVenus and WT mVenus/chlorophyll fluorescence in replete (36 µM phosphate), deplete (1.8 µM phosphate) and resupply (1.8 µM day 4 cells resupplied with 36 µM Pi) without RuR. **D**) As in C, except RuR was applied to all treatments on day 4, just prior to phosphate resupply in the resupply treatment. Data in C and D are mean ± standard error (n=3).

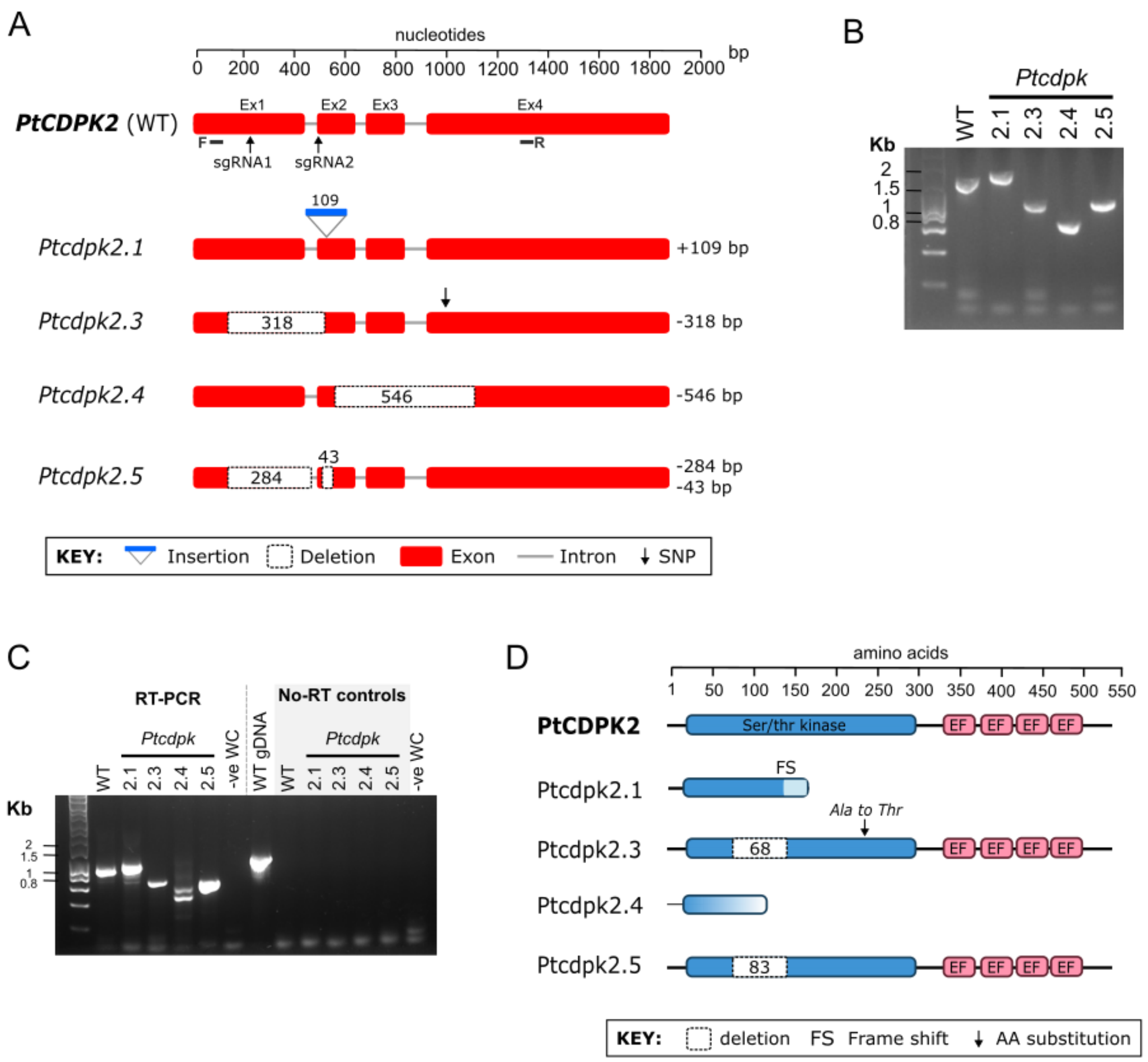


**Figure S4. Generation of biallelic *Ptcdpk2* mutants in *Phaeodactylum tricornutum* using CRISPR-Cas9 gene editing. A)** Nucleotide sequence of WT *PtCDPK2* (Phatr2_21006) and *Ptcdpk2* mutant lines. Exons are red, and grey lines are introns. Black arrows indicate specific single guide RNA locations. Primer binding sites for Keh 409F (F) and Keh 410R (R) used for PCR screening and analysis are also indicated. Sizes of indels are indicated to the right of the gene alignment, as identified through band-shift gel electrophoresis followed by Sanger sequencing. A single nucleotide polymorphism (SNP) is also indicated in *Ptcdpk2.3*. **B)** PCR amplified DNA fragments of the target site in wildtype (WT) and *Ptcdpk2* mutant lines using Keh 409F (F) and Keh 410R (R). Expected product size for WT was 1166 bp. The gel was stained with Gel Red (Biotium), which in our hands can lead to discrepancies between ladder sizes and expected product sizes. Hence all bands were sequenced via Sanger sequencing to confirm their size and nucleotide sequences (**given in Supplementary Information 3-6**). Sequencing data revealed no evidence of ambiguous nucleotide sequences, indicating identical indels on both alleles. **C)** PCR amplification of cDNA from WT and *Ptcdpk2* mutant lines using primers Keh 409F and Keh 410R via reverse transcriptase (RT)-PCR (**Supplementary Methods**). Negative no template water controls (-ve WC), as well as no-RT controls using RNA samples but with *Taq* polymerase, without reverse transcriptase are indicated. No-RT controls were further validated using genomic DNA extracted from WT *P. tricornutum* (WT gDNA). PCR products were analysed by agarose gel electrophoresis (1% agarose, stained with GelRed). **D)** Predicted protein structure deduced from the RT-PCR analysis (**Supplementary Methods**), indicating positions of deletions, substitutions, frameshifts and/or truncations in mutants compared to WT. The SNP identified in the DNA (and corresponding cDNA sequence) of *Ptcdpk2.3,* would cause an alanine (Ala) to threonine (Thr) substitution in a conserved region of the protein, as indicated.
