## Supplemental Information for "A plasma membrane Ca^2+^-dependent protein kinase PtCDPK2 promotes phosphorus starvation resilience in *Phaeodactylum tricornutum*"

**Supplementary information 1:** Gene model for *PtCDPK2* (protein id. 21006), from *P. tricornutum* genome v3. To generate the PtCDPK2-mVenus construct this sequence was synthesised, excluding the TGA stop codon but including flanking restrictions site sequences for PstI (GGGCTGCAG) and StuI (AGGCCT). This DNA fragment was synthesised and cloned by Genscript (GenScript, Piscataway, NJ) into the pPha-T1-mVenus vector (a derivative of the pPHA-T1 (accession AF219942) adapted as described by (Helliwell et al., 2019) containing a codon optimised mVenus gene (AJN91098.1).

XX = Upstream untranslated region
XX = Exon
XX = Intron

>PtCDPK21006_ gene model
GTCTACAAAAGCATGTCTTGTAAGCTAATCCATCCTTTATAGATCAGGATTGGAATAACCTGCCAAAA
ACCTACTACCTATAGAGTTCAATTCTGGGTCAGCTTCTTTCAATGGTGACAGTAGTTCCGTCACATAA
AGCGGTGGCCACTTGTTTCTCCCTTCGTGGTGTCCGTGACTCTCAATTGTGAAAAACCCTCACTCTCT
GGTGAAAAGTGAAGAAAACCAGGGTAGAGTAGGAACTAGCAGGTTTCTTGCTGACAGCGAATCCAT
TCTTACGCTCATGTAGAGATGTGCAGTCTGAGGGGTTGGTCAAGAGAATGTTGCGGGGCCATGTTA
CACTCGTGGCGGTGTGTAGCGAATTTGTTTCCACTCATTGTAGAGAAAAACAACAAAGATAAAAAG
GAAAATCCACTATATGGGATACTACGATTCGCAGTCTTTTGAAGACCCGGAAATGAGAATCTCAGCG
TCCGTTGAGCGAATCTAACGTTAGGTTTCGAAATTTTGAAGTGATCGCGCCGCGTAGGAATCAGCAA
AAAACAGCCGAAGAAGATCCAAGCTGCCGACGACCGCTCGGCTCGCACAAGGAACGACCGAGTTCA
AGACAGAACCAGCAGCCAGAATGTTATGTAGACCTTGTTGTGACGAAGACAGCGACAAATCTGCTT
CTCGTAGCCTGTTCACACCACATATTCAAGCTATGGACGAACTTCCTATTACCGACCCTTCCATCCATG
CCAACCTCCTACTCGAATCGAAGGAGGATGTCTTCAAAAAATATTCCGTCGTCCAAGTGCTCGGTAA
TGGATCAATGGGAACCGTTTCCAAGGTCAAAATTAAGAAGCACAAGGTCGGGGGAAGCGCCTTTCA
GCCGAAATCCAAGGGAATTTTTGGCTTTTTGAAGAAACAGAACAACAAAAGGAAGGAAGGTGAGA
CCAGAGAACACAATAGTCAGGACTATATATACGCACTCAAGTCCATCATTCTGGATCGGGTCTCTTCT
GTCTTCCTGGACGAGCTCCGTAACGAAATTCTTATCCTTAGATCATTGGATCATCCCAATATTGTCAA
AGCGCACGAGGTTTACTACACGAGGAAGCAGATTTATCTCGGTGCGTGATGAGGAATGTGAAACGT
GATTGGTCAGAAGTCTGAACTTTTTCTCTTTGGAAAGGAGACTAACGCATTTCCGTCATCGTTTTCTT
ATATCCGTGTGCTGTTTCTGGAAGTATTGGAGTTGTGTGATGGCGGAGACCTTTATACCAGGTCGCC
TTACAGTGAAAGGGAATCGGCAAGGATTCTGCAACAAATATTGTCGGCAGTGCGGTACATGCATGG
TACGCTATACCGATACGCATATAGAATTCCCGCAGCAAACTTTGGGAAGCTAACAATTCTTGTTCTGC
TTTGTCTAGATCACGGAATTGTTCATCGGGATCTCAAGTTCGAGAATATCATGTTTGAGAACAATAG
CCCCAGTGCTCGGTAGGTTACTGATCAACTCTCGAGGCTACACGTAGATGAAGCGCTCTTACAGTCA
ATAATGTTCTATTCATTTTGCACAGAGTCAAAATTATAGATTTTGGATTGTCTAAAAAGTTCCTTGGC
AAACCGTCGTACATGACCGAACGCGTTGGTACCGTCTATACGATGGCCCCGCAAGTCCTGCAAGGA
GTCTACTCATCGCAAGCTGATCTTTGGTCCGCTGGAGTGATAGCCTACATGCTGTTATCGGCTTCAAA
GCCTTTTTATCACAAACGACGGCGCAAGATGATTGACCAAATCATGAGGGCCGACTTCGGATATAAT
GCACCGGTCTGGAAGCAAATATCAGAAAGTGCGCAAGATTTTGTAAGTCGATTACTAGTGGTGGAT
CCAAAGAAAAGACTGAATGCAGAAAAAGCATTGGACCATTCTTGGATTGTGAATCGCGAACGCTTG
CCAGATGAGACACCATCCGAGGATTTGTTGGCCGCTGTCGATGATTGCCTCGTGAATTATCGACAAA
CGTCGGAGCTGAAAAAGCTAGCTTTAAACATGATCGCCCATCGTTCTACCGCGGAAGAGATCATGCA
ACTTCGGAAAGTTTTTGACAGCTACGACACCTCGAATGATGGAATTATTACATTTGATGAATTCAAAG
CAGCTTTGCACAAAATGAAATATCCGGATGAGATTGTACAGGAAGTTTTTAGCAGTATTGATGTCAA
CCGAAATGGCCATATACAGTACACGGAATTCATTGCATCGACCGTCTTGGCACAGGGACATATCGCA

GAGGATCGGGTCGCAGTAGCTTTCGATCGCTTGGACTCTGATGACACCGGCTTTATTTCCAAGAAGA
ACTTGCAAAACGCATTGGGCAAGGAATACACTCCAGAACTCGTCGAAAATATAATGGAAGAAGTTG
ACAAAGATAGGGATGGCAAAATATCATATACCGAGTTTCTGCAATACTTTCGGAAGGAAACGAGCA
ATCTGGCCGACAGGGCCTCTCTTTTAGAGCAACAGTCGTCACATGTAAGCGAACACGGTCTGGTCGG
TTTGGACGCCAAGATTCCTGGAGGACCGTACGATCCTAACCGAACT**TGA**

**Supplementary information 2:** Description and features of DNA fragment synthesised and cloned into pKSdiaCas9 (Nymark et al., 2016) using restriction site/enzyme BsaI. The resulting plasmid was subsequently co-transformed into *P. tricornutum* with pPHAT1 (accession no.: AF219942) conferring resistance to zeocin to generate *Ptcdpk2* mutants described in this paper.

XX = BsaI bind
XX = BsaI cut
XX = sgRNA1_Pt21006
XX = sgRNA2_Pt21006
XX = sgRNA Cas9bind
XX = Pt U6Promoter

>CDPK 21006_dual sgRNA cassette

GGTCTCATCGAGAAACAGAACAACAAAAGGAGTTTTAGAGCTAGAAATAGCAAGTTAAAATAAGGC
TAGTCCGTTATCAACTTGAAAAAGTGGCACCGAGTCGGTGCTTTTTTTCTAGACCCAGCTTTCTTGTA
CAAAGTTGGCATTACGCTTTACGAATTCCCATGGGGAGGTTGGCTCGGAAGTTGGTGTTGACGGTG
AGCTGGAAATTGGTTGTCGGTCACTGCTAGCGAGAAGAAAACGGAGGACAGAAGGAAGTGAAACT
CGGTTCGTTCTCGACAGCCTCACTGTCAATATGCTCATTTTCAATCCTTAGCGCTTTTAATGTCGAATT
GACGGTAAATTGAATAGGATCTATAATATCTACAAGGTACTTTGACACGCCAAGTATTCATTGTTAGT
CAACAATATTTTAGAGCTTTATAAGGTCAAAAAACACCTTCAAAGTCGAGGAAGTATTGGAGTTGTG
TGAGTTTAGAGACC

**Supplementary information 3:** DNA sequence of *Ptcdpk2.1*

Insertion
XX = Exon
XX = Intron
**XX**= Primer binding sites for KEH_409F and KEH_410R

*>Ptcdpk2.1*
ATGGACGAACTTCCTATTACCGACCCTTCCATCCATGCCAACCTCCTACTCGAATCGAAGGAGGATGT
CTTCAAAAAATATTCCGT**CGTCCAAGTGCTCGGTAATG**GATCAATGGGAACCGTTTCCAAGGTCAAA
ATTAAGAAGCACAAGGTCGGGGGAAGCGCCTTTCAGCCGAAATCCAAGGGAATTTTTGGCTTTTTG
AAGAAACAGAACAACAAAAGGAAGGAAGGTGAGACCAGAGAACACAATAGTCAGGACTATATATA
CGCACTCAAGTCCATCATTCTGGATCGGGTCTCTTCTGTCTTCCTGGACGAGCTCCGTAACGAAATTC
TTATCCTTAGATCATTGGATCATCCCAATATTGTCAAAGCGCACGAGGTTTACTACACGAGGAAGCA
GATTTATCTCGGTGCGTGATGAGGAATGTGAAACGTGATTGGTCAGAAGTCTGAACTTTTTCTCTTT
GGAAAGGAGACTAACGCATTTCCGTCATCGTTTTCTTATATCCGTGTGCTGTTTCTGGAAGTATTGGA
GTTGTGTGAGAAAACTCACCTCTGCCTAGCCTACACGACGCTGAAGCTATGTGTCTCGAGAAAACTC
ATCCTGTGCCTTCCTCACTTTCTGCTAGTCCACTCCAACAATACGGCGTGATGGCGGAGACCTTTATA
CCAGGTCGCCTTACAGTGA**AAGGG**AATCGGCAAGGATTCTGCAACAAATATTGTCGGCAGTGCGGT
ACATGCATGGTACGCTATACCGATACGCATATAGAATTCCCGCAGCAAACTTTGGGAAGCTAACAAT
TCTTGTTCTGCTTTGTCTAGATCACGGAATTGTTCATCGGGATCTCAAGTTCGAGAATATCATGTTTG
AGAACAATAGCCCCAGTGCTCGGTAGGTTACTGATCAACTCTCGAGGCTACACGTAGATGAAGCGCT
CTTACAGTCAATAATGTTCTATTCATTTTGCACAGAGTCAAAATTATAGATTTTGGATTGTCTAAAAA
GTTCCTTGGCAAACCGTCGTACATGACCGAACGCGTTGGTACCGTCTATACGATGGCCCCGCAAGTC
CTGCAAGGAGTCTACTCATCGCAAGCTGATCTTTGGTCCGCTGGAGTGATAGCCTACATGCTGTTAT
CGGCTTCAAAGCCTTTTTATCACAAACGACGGCGCAAGATGATTGACCAAATCATGAGGGCCGACTT
CGGATATAATGCACCGGTCTGGAAGCAAATATCAGAAAGTGCGCAAGATTTTGTAAGTCGATTACTA
GTGGTGGATCCAAAGAAAAGACTGAATGCAGAAAAAGCATTGGACCATTCTTGGATTGTGAATCGC
**GAACGCTTGCCAGATGAGAC**ACCATCCGAGGATTTGTTGGCCGCTGTCGATGATTGCCTCGTGAATT
ATCGACAAACGTCGGAGCTGAAAAAGCTAGCTTTAAACATGATCGCCCATCGTTCTACCGCGGAAGA
GATCATGCAACTTCGGAAAGTTTTTGACAGCTACGACACCTCGAATGATGGAATTATTACATTTGAT
GAATTCAAAGCAGCTTTGCACAAAATGAAATATCCGGATGAGATTGTACAGGAAGTTTTTAGCAGTA
TTGATGTCAACCGAAATGGCCATATACAGTACACGGAATTCATTGCATCGACCGTCTTGGCACAGGG
ACATATCGCAGAGGATCGGGTCGCAGTAGCTTTCGATCGCTTGGACTCTGATGACACCGGCTTTATT
TCCAAGAAGAACTTGCAAAACGCATTGGGCAAGGAATACACTCCAGAACTCGTCGAAAATATAATG
GAAGAAGTTGACAAAGATAGGGATGGCAAAATATCATATACCGAGTTTCTGCAATACTTTCGGAAG
GAAACGAGCAATCTGGCCGACAGGGCCTCTCTTTTAGAGCAACAGTCGTCACATGTAAGCGAACAC
GGTCTGGTCGGTTTGGACGCCAAGATTCCTGGAGGACCGTACGATCCTAACCGAACT**TGA**

**Supplementary information 4:** DNA sequence of *Ptcdpk2.3*

Region deleted
XX = Exon
XX = Intron
**XX**= Primer binding sites for KEH_409F and KEH_410R
X= single nucleotide polymorphism confirmed at DNA and cDNA level, causing a change from Ala to Thr in the protein sequence.

*>Ptcdpk2.3*

ATGGACGAACTTCCTATTACCGACCCTTCCATCCATGCCAACCTCCTACTCGAATCGAAGGAGGATGT
CTTCAAAAAATATTCCGT**CGTCCAAGTGCTCGGTAATG**GATCAATGGGAACCGTTTCCAAGGTCAAA
ATTAAGAAGCACAAGGTCGGGGGAAGCGCCTTTCAGCCGAAATCCAAGGGAATTTTTGGCTTTTTG
AAGAAACAGAACAACAAAAGGAAGGAAGGTGAGACCAGAGAACACAATAGTCAGGACTATATATA
CGCACTCAAGTCCATCATTCTGGATCGGGTCTCTTCTGTCTTCCTGGACGAGCTCCGTAACGAAATTC
TTATCCTTAGATCATTGGATCATCCCAATATTGTCAAAGCGCACGAGGTTTACTACACGAGGAAGCA
GATTTATCTCGGTGCGTGATGAGGAATGTGAAACGTGATTGGTCAGAAGTCTGAACTTTTTCTCTTT
GGAAAGGAGACTAACGCATTTCCGTCATCGTTTTCTTATATCCGTGTGCTGTTTCTGGAAGTATTGGA
GTTGTGTGATGGCGGAGACCTTTATACCAGGTCGCCTTACAGTGA**AAGGG**AATCGGCAAGGATTCT
GCAACAAATATTGTCGGCAGTGCGGTACATGCATGGTACGCTATACCGATACGCATATAGAATTCCC
GCAGCAAACTTTGGGAAGCTAACAATTCTTGTTCTGCTTTGTCTAGATCACGGAATTGTTCATCGGGA
TCTCAAGTTCGAGAATATCATGTTTGAGAACAATAGCCCCAGTGCTCGGTAGGTTACTGATCAACTCT
CGAGGCTACACGTAGATGAAGCGCTCTTACAGTCAATAATGTTCTATTCATTTTGCACAGAGTCAAA
ATTATAGATTTTGGATTGTCTAAAAAGTTCCTTGGCAAACCGTCGTACATGACCGAACGCGTTGGTA
CCGTCTATACGATGGCCCCGCAAGTCCTGCAAGGAGTCTACTCATCGCAAGCTGATCTTTGGTCCACT
GGAGTGATAGCCTACATGCTGTTATCGGCTTCAAAGCCTTTTTATCACAAACGACGGCGCAAGATGA
TTGACCAAATCATGAGGGCCGACTTCGGATATAATGCACCGGTCTGGAAGCAAATATCAGAAAGTG
CGCAAGATTTTGTAAGTCGATTACTAGTGGTGGATCCAAAGAAAAGACTGAATGCAGAAAAAGCAT
TGGACCATTCTTGGATTGTGAATCGC**GAACGCTTGCCAGATGAGAC**ACCATCCGAGGATTTGTTGG
CCGCTGTCGATGATTGCCTCGTGAATTATCGACAAACGTCGGAGCTGAAAAAGCTAGCTTTAAACAT
GATCGCCCATCGTTCTACCGCGGAAGAGATCATGCAACTTCGGAAAGTTTTTGACAGCTACGACACC
TCGAATGATGGAATTATTACATTTGATGAATTCAAAGCAGCTTTGCACAAAATGAAATATCCGGATG
AGATTGTACAGGAAGTTTTTAGCAGTATTGATGTCAACCGAAATGGCCATATACAGTACACGGAATT
CATTGCATCGACCGTCTTGGCACAGGGACATATCGCAGAGGATCGGGTCGCAGTAGCTTTCGATCG
CTTGGACTCTGATGACACCGGCTTTATTTCCAAGAAGAACTTGCAAAACGCATTGGGCAAGGAATAC
ACTCCAGAACTCGTCGAAAATATAATGGAAGAAGTTGACAAAGATAGGGATGGCAAAATATCATAT
ACCGAGTTTCTGCAATACTTTCGGAAGGAAACGAGCAATCTGGCCGACAGGGCCTCTCTTTTAGAGC
AACAGTCGTCACATGTAAGCGAACACGGTCTGGTCGGTTTGGACGCCAAGATTCCTGGAGGACCGT
ACGATCCTAACCGAACT**TGA**

**Supplementary information 5:** DNA sequence of *Ptcdpk2.4*

Region deleted
XX = Exon
XX = Intron
**XX**= Primer binding sites for KEH_409F and KEH_410R

*>Ptcdpk2.4*
ATGGACGAACTTCCTATTACCGACCCTTCCATCCATGCCAACCTCCTACTCGAATCGAAGGAGGATGT
CTTCAAAAAATATTCCGT**CGTCCAAGTGCTCGGTAATG**GATCAATGGGAACCGTTTCCAAGGTCAAA
ATTAAGAAGCACAAGGTCGGGGGAAGCGCCTTTCAGCCGAAATCCAAGGGAATTTTTGGCTTTTTG
AAGAAACAGAACAACAAAAGGAAGGAAGGTGAGACCAGAGAACACAATAGTCAGGACTATATATA
CGCACTCAAGTCCATCATTCTGGATCGGGTCTCTTCTGTCTTCCTGGACGAGCTCCGTAACGAAATTC
TTATCCTTAGATCATTGGATCATCCCAATATTGTCAAAGCGCACGAGGTTTACTACACGAGGAAGCA
GATTTATCTCGGTGCGTGATGAGGAATGTGAAACGTGATTGGTCAGAAGTCTGAACTTTTTCTCTTT
GGAAAGGAGACTAACGCATTTCCGTCATCGTTTTCTTATATCCGTGTGCTGTTTCTGGAAGTATTGGA
GTTGTGTGATGGCGGAGACCTTTATACCAGGTCGCCTTACAGTGA**AAGGG**AATCGGCAAGGATTCT
GCAACAAATATTGTCGGCAGTGCGGTACATGCATGGTACGCTATACCGATACGCATATAGAATTCCC
GCAGCAAACTTTGGGAAGCTAACAATTCTTGTTCTGCTTTGTCTAGATCACGGAATTGTTCATCGGGA
TCTCAAGTTCGAGAATATCATGTTTGAGAACAATAGCCCCAGTGCTCGGTAGGTTACTGATCAACTCT
CGAGGCTACACGTAGATGAAGCGCTCTTACAGTCAATAATGTTCTATTCATTTTGCACAGAGTCAAA
ATTATAGATTTTGGATTGTCTAAAAAGTTCCTTGGCAAACCGTCGTACATGACCGAACGCGTTGGTA
CCGTCTATACGATGGCCCCGCAAGTCCTGCAAGGAGTCTACTCATCGCAAGCTGATCTTTGGTCCGC
TGGAGTGATAGCCTACATGCTGTTATCGGCTTCAAAGCCTTTTTATCACAAACGACGGCGCAAGATG
ATTGACCAAATCATGAGGGCCGACTTCGGATATAATGCACCGGTCTGGAAGCAAATATCAGAAAGT
GCGCAAGATTTTGTAAGTCGATTACTAGTGGTGGATCCAAAGAAAAGACTGAATGCAGAAAAAGCA
TTGGACCATTCTTGGATTGTGAATCGC**GAACGCTTGCCAGATGAGAC**ACCATCCGAGGATTTGTTGG
CCGCTGTCGATGATTGCCTCGTGAATTATCGACAAACGTCGGAGCTGAAAAAGCTAGCTTTAAACAT
GATCGCCCATCGTTCTACCGCGGAAGAGATCATGCAACTTCGGAAAGTTTTTGACAGCTACGACACC
TCGAATGATGGAATTATTACATTTGATGAATTCAAAGCAGCTTTGCACAAAATGAAATATCCGGATG
AGATTGTACAGGAAGTTTTTAGCAGTATTGATGTCAACCGAAATGGCCATATACAGTACACGGAATT
CATTGCATCGACCGTCTTGGCACAGGGACATATCGCAGAGGATCGGGTCGCAGTAGCTTTCGATCG
CTTGGACTCTGATGACACCGGCTTTATTTCCAAGAAGAACTTGCAAAACGCATTGGGCAAGGAATAC
ACTCCAGAACTCGTCGAAAATATAATGGAAGAAGTTGACAAAGATAGGGATGGCAAAATATCATAT
ACCGAGTTTCTGCAATACTTTCGGAAGGAAACGAGCAATCTGGCCGACAGGGCCTCTCTTTTAGAGC
AACAGTCGTCACATGTAAGCGAACACGGTCTGGTCGGTTTGGACGCCAAGATTCCTGGAGGACCGT
ACGATCCTAACCGAACT**TGA**

**Supplementary information 6:** DNA sequence of *Ptcdpk2.5*

Deleted region 1
Deleted region 2
XX = Exon
XX = Intron
**XX**= Primer binding sites for KEH_409F and KEH_410R

*>Ptcdpk2.5*
ATGGACGAACTTCCTATTACCGACCCTTCCATCCATGCCAACCTCCTACTCGAATCGAAGGAGGATGT
CTTCAAAAAATATTCCGT**CGTCCAAGTGCTCGGTAATG**GATCAATGGGAACCGTTTCCAAGGTCAAA
ATTAAGAAGCACAAGGTCGGGGAAGCGCCTTTCAGCCGAAATCCAAGGGAATTTTTGGCTTTTTG
AAGAAACAGAACAACAAAAGGAAGGAAGGTGAGACCAGAGAACACAATAGTCAGGACTATATATA
CGCACTCAAGTCCATCATTCTGGATCGGGTCTCTTCTGTCTTCCTGGACGAGCTCCGTAACGAAATTC
TTATCCTTAGATCATTGGATCATCCCAATATTGTCAAAGCGCACGAGGTTTACTACACGAGGAAGCA
GATTTATCTCGGTGCGTGATGAGGAATGTGAAACGTGATTGGTCAGAAGTCTGAACTTTTTCTCTTT
GGAAAGGAGACTAACGCATTTCCGTCATCGTTTTCTTATATCCGTGTGCTGTTTCTGGAAGTATTGGA
GTTGTGTGATGGCGGAGACCTTTATACCAGGTCGCCTTACAGTGA**AAGGG**AATCGGCAAGGATTCT
GCAACAAATATTGTCGGCAGTGCGGTACATGCATGGTACGCTATACCGATACGCATATAGAATTCCC
GCAGCAAACTTTGGGAAGCTAACAATTCTTGTTCTGCTTTGTCTAGATCACGGAATTGTTCATCGGGA
TCTCAAGTTCGAGAATATCATGTTTGAGAACAATAGCCCCAGTGCTCGGTAGGTTACTGATCAACTCT
CGAGGCTACACGTAGATGAAGCGCTCTTACAGTCAATAATGTTCTATTCATTTTGCACAGAGTCAAA
ATTATAGATTTTGGATTGTCTAAAAAGTTCCTTGGCAAACCGTCGTACATGACCGAACGCGTTGGTA
CCGTCTATACGATGGCCCCGCAAGTCCTGCAAGGAGTCTACTCATCGCAAGCTGATCTTTGGTCCGC
TGGAGTGATAGCCTACATGCTGTTATCGGCTTCAAAGCCTTTTTATCACAAACGACGGCGCAAGATG
ATTGACCAAATCATGAGGGCCGACTTCGGATATAATGCACCGGTCTGGAAGCAAATATCAGAAAGT
GCGCAAGATTTTGTAAGTCGATTACTAGTGGTGGATCCAAAGAAAAGACTGAATGCAGAAAAAGCA
TTGGACCATTCTTGGATTGTGAATCGC**GAACGCTTGCCAGATGAGAC**ACCATCCGAGGATTTGTTGG
CCGCTGTCGATGATTGCCTCGTGAATTATCGACAAACGTCGGAGCTGAAAAAGCTAGCTTTAAACAT
GATCGCCCATCGTTCTACCGCGGAAGAGATCATGCAACTTCGGAAAGTTTTTGACAGCTACGACACC
TCGAATGATGGAATTATTACATTTGATGAATTCAAAGCAGCTTTGCACAAAATGAAATATCCGGATG
AGATTGTACAGGAAGTTTTTAGCAGTATTGATGTCAACCGAAATGGCCATATACAGTACACGGAATT
CATTGCATCGACCGTCTTGGCACAGGGACATATCGCAGAGGATCGGGTCGCAGTAGCTTTCGATCG
CTTGGACTCTGATGACACCGGCTTTATTTCCAAGAAGAACTTGCAAAACGCATTGGGCAAGGAATAC
ACTCCAGAACTCGTCGAAAATATAATGGAAGAAGTTGACAAAGATAGGGATGGCAAAATATCATAT
ACCGAGTTTCTGCAATACTTTCGGAAGGAAACGAGCAATCTGGCCGACAGGGCCTCTCTTTTAGAGC
AACAGTCGTCACATGTAAGCGAACACGGTCTGGTCGGTTTGGACGCCAAGATTCCTGGAGGACCGT
ACGATCCTAACCGAACT**TGA**
