## Supplemental Methods for "A plasma membrane Ca^2+^-dependent protein kinase PtCDPK2 promotes phosphorus starvation resilience in *Phaeodactylum tricornutum*"

**Supplementary Methods**

RNA extraction and RT-PCR of *PtCDPK2* gene in *Ptcdpk2* mutants

To further clarify resulting coding sequences of WT and *Ptcdpk2* mutants, RT-PCR was performed. Strains were grown in 100 ml f/2 medium with 1.8 µM phosphate, without silicate and vitamins (to an $OD_{730}$ of approximately 0.1, and 1–2 x $10^6$ cells/ml). Cells were harvested in 50 ml aliquots by centrifugation (4,000 g, 10 mins). Cell pellets were weighed then snap frozen in liquid nitrogen for storage at -80°C prior to processing. RNA was extracted from cell pellets (≤100 mg wet weight) using the RNeasy Plant Mini Kit (Qiagen). Cells were thawed on ice in 450 µl RLT (a lysis buffer containing guanidine isothiocycanate, Qiagen RNeasy kit), with additional 1% v/v Beta-mercaptoethanol and approximately 300 µl (equivalent volume) of glass beads (0.1–1.5 mm, autoclaved then pre-soaked in RLT buffer overnight). Cells were lysed by vortexing (5 x 10 secs, alternated with 10 secs chilling on ice). Cell debris and un-lysed cells were removed by centrifugation (200 g, 10 secs, then 17,000 g, for 1 min). RNA was processed according to the manufacturer's instructions, except the column was washed 3 times with RPE buffer. DNA contamination was eliminated using a Turbo DNA-*free* Kit (Invitrogen, as in the manufacturer's instructions) and RNA quantified by NanoDrop (Thermo Scientific) and frozen in aliquots at -80°C.

RT-PCR was performed on the RNA samples (~20 ng) using a One*Taq* One-Step RT-PCR kit (NEB, according to manufacturer's instructions), with primers keh409F CGTCCAAGTGCTCGGTAATG and keh410R GTCTCATCTGGCAAGCGTTC. RT-PCR cycling conditions: 48°C for 15 mins (cDNA synthesis step), followed immediately by 94°C 60 secs, 40 x (94°C 15 secs, 53°C 30 secs, 68°C 60 secs), and 68°C 5 minutes. Negative controls, using the same primers but PCR rather than RT-PCR conditions (standard *Taq* instead of the One*Taq* RT-PCR enzyme mix, and no initial 15 minutes at 48°C), were conducted to confirm lack of DNA contamination of RNA samples. As a further control, purified genomic DNA was amplified by these PCR conditions. RT-PCR products and PCR controls were analysed by gel electrophoresis (1% agarose, stained with GelRed), and sequenced (Source BioScience, via Sanger sequencing with primers keh409F and keh410R).

For mutant *Ptcdpk2.4,* RT-PCR yielded a double band despite no evidence of DNA contamination in the no-RT controls (**Figure S4C**), and so further analysis was required. Each band was gel isolated and cleaned (Nucleospin Gel & PCR Clean-up, Macherey-Nagel) for sequencing. In addition, the mixed RT-PCR product was cloned into the pCR 2.1 vector (TA cloning kit, Invitrogen) with blue/white selection (LB Amp, XGal, IPTG plates). Colony PCRs of white colonies (nine in total), using GoTaq (Promega) and universal M13 primers (M13F-20 GTAAAACGACGGCCAG, M13R CAGGAAACAGCTATGACC) were conducted using PCR cycling conditions: 95°C 3 mins, 31 x (95°C 30 sec, 46°C 30 sec, 72°C 35 sec), and 72°C 5 min. Clones with inserts were sent for sequencing with the M13 primers (Source BioScience). All sequences were checked for ambiguities, trimmed and aligned in BioEdit (RRID:SCR_007361) and translated with ExPASy (Swiss Bioinformatics Resource Portal). Sequencing analysis revealed sequence variation, including alternative splicing of intron 1. In addition, there was a small variable region (with a single nucleotide polymorphism, a 1 bp insert, and deletions of 8 bp or 21 bp observed). Translation of *Ptcdpk2.4* cDNA sequences indicated that all variants analysed would produce severely truncated proteins (74 – 141 amino acids, compared to the full length of 538) disrupting protein function. The sequence variations observed are likely due to re-editing in this mutant line, as has been previously reported for *P. tricornutum* mutants over extended growth maintenance periods (Sharma et al., 2018).
